# Vision-dependent and -independent molecular maturation of mouse retinal ganglion cells

**DOI:** 10.1101/2022.04.20.488897

**Authors:** Irene E. Whitney, Salwan Butrus, Michael A. Dyer, Fred Rieke, Joshua R. Sanes, Karthik Shekhar

## Abstract

The development and connectivity of retinal ganglion cells (RGCs), the retina’s sole output neurons, are patterned by activity-independent transcriptional programs and activity-dependent remodeling. To inventory the molecular correlates of these influences, we applied high-throughput single-cell RNA sequencing (scRNA-seq) to mouse RGCs at six embryonic and postnatal ages. We identified temporally regulated modules of genes that correlate with, and likely regulate, multiple phases of RGC development, ranging from differentiation and axon guidance to synaptic recognition and refinement. Some of these genes are expressed broadly while others, including key transcription factors and recognition molecules, are selectively expressed by one or a few of the 45 transcriptomically distinct types defined previously in adult mice. Next, we used these results as a foundation to analyze the transcriptomes of RGCs in mice lacking visual experience due to dark rearing from birth or to mutations that ablate either bipolar or photoreceptor cells. 98.5% of visually deprived (VD) RGCs could be unequivocally assigned to a single RGC type based on their transcriptional profiles, demonstrating that visual activity is dispensable for acquisition and maintenance of RGC type identity. However, visual deprivation significantly reduced the transcriptomic distinctions among RGC types, implying that activity is required for complete RGC maturation or maintenance. Consistent with this notion, transcriptomic alternations in VD RGCs significantly overlapped with gene modules found in developing RGCs. Our results provide a resource for mechanistic analyses of RGC differentiation and maturation, and for investigating the role of activity in these processes.

## INTRODUCTION

Retinal ganglion cells (RGCs) are the sole output neurons of the retina (Figure 1A). They receive and integrate visual information from photoreceptors via interneurons and project axons to the rest of the brain. In mice, RGCs can be divided into ∼45 discrete types that are distinguishable by their morphological, physiological and molecular properties (Baden et al., 2016; Bae et al., 2018; Goetz et al., 2021; Rheaume et al., 2018; Tran et al., 2019). Many of these types respond selectively to particular visual features such as motion or edges, thereby parcellating visual information into parallel channels that are transmitted to numerous central brain targets (Dhande et al., 2015; Martersteck et al., 2017). The structure, function and development of RGCs have been extensively studied, making them among the best-characterized of vertebrate central neuronal classes (Sanes and Masland, 2015). Consequently, studying the genesis, specification and differentiation of RGCs can not only help elucidate principles that govern the development of the visual system, but also inform our understanding of neural development generally.

**Figure 1.**
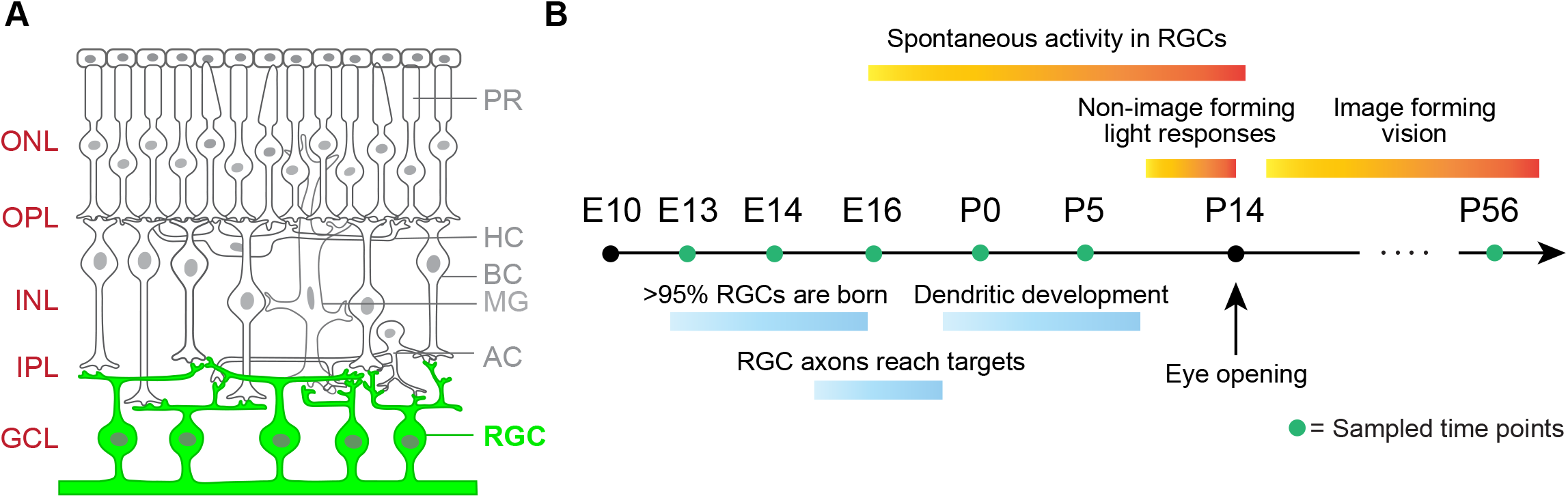
Retinal ganglion cell (RGC) development in mice. A. Sketch of a cross-section of the retina, highlighting retinal ganglion cells (RGCs) located in the ganglion cell layer (GCL). PR, photoreceptors; HC, horizontal cells; BC, bipolar cells; MG, Müller glia; AC, amacrine cells; ONL, outer nuclear layer; OPL, outer plexiform layer; INL, inner nuclear layer; IPL, inner plexiform layer, B. Timeline of RGC development in mice highlighting key developmental milestones. Green dots indicate ages considered in this study.

The development of RGCs and their integration into circuits are orchestrated by a combination of activity-independent and -dependent influences. Nowhere is this more evident than in the retinotectal projection (retinocollicular in mammals). Sperry hypothesized that RGCs and their tectal targets bear graded “chemoaffinity” labels that lead to the orderly retinotopic mapping of the dorso-ventral and anterior-posterior axes of the retina onto those of the tectum (Sperry, 1963). In a long series of experiments, Bonhoeffer and colleagues devised *ex vivo* assays that recapitulated essential features of retinotopic matching (Bonhoeffer and Huf, 1985; Walter et al., 1987a; Walter et al., 1987b), leading eventually to the demonstration that opposing gradients of Eph kinases and their ligands, the ephrins, underlie the establishment of this mapping (Cheng et al., 1995; Drescher et al., 1995; Feldheim and O’Leary, 2010; Stahl et al., 1990a; Stahl et al., 1990b). On the other hand, the “hard-wired” retinotopic map is somewhat imprecise, and is extensively refined by activity-dependent mechanisms, leading to the more accurate topography seen in adulthood (Cang et al., 2005; Feldheim and O’Leary, 2010; McLaughlin et al., 2003; Schmidt and Buzzard, 1993).

Other features of RGCs are also shaped by an interplay of activity-dependent and -independent influences. For example, the segregation of RGC axons into discrete laminae in the lateral geniculate nucleus requires initial molecular recognition followed by activity-dependent refinement, although in this case, little is known about the molecular basis of the initial phases (Hooks and Chen, 2007; 2008; Katz and Shatz, 1996). Within the retina, the dendritic arbors of specific RGC types appear to be shaped predominantly by intrinsic transcriptional programs (Liu et al., 2018; Peng et al., 2017) that also specify the interneuronal types from which they receive synapses (Duan et al., 2014; Krishnaswamy et al., 2015; Sanes and Zipursky, 2020). However, visual deprivation and/or decreased activity affect both dendritic morphology (Bodnarenko et al., 1995; Chalupa and Gunhan, 2004; El-Quessny et al., 2020; Tian and Copenhagen, 2003) and patterns of synaptic input (Arroyo and Feller, 2016; Kerschensteiner et al., 2009; Morgan et al., 2011; Okawa et al., 2014).

Interactions of activity-independent and -dependent influences occur dynamically and over a protracted period. The transcriptomes of RGCs change dramatically as they diversify, develop and mature (see **Results**). Likewise, activity arises from multiple sources that change during development. In late embryonic and early postnatal life in rodents, RGC activity is “spontaneous”, arising from intrinsic biophysical properties of neighboring interneurons and their connections to each other and to RGCs (Arroyo and Feller, 2016; Wong, 1999). Later, but prior to eye-opening, light-dependent activity affects development by non-image forming activation of rod and cone photoreceptors (Tiriac et al., 2018). Later still, conventional visual input affects maturation of RGC dendritic and axonal arbors as well as their maintenance (see above). Finally, both pre- and postnatally, activation of melanopsin-containing intrinsically photosensitive RGCs (ipRGCs) affects development of retinal vasculature and patterns of spontaneous activity (Kirkby and Feller, 2013).

Elucidating the mechanisms that regulate RGC development and the ways in which activity-independent and -dependent influences interact would be greatly aided by comprehensive characterization of gene expression in developing RGCs. Our aim in this study is to provide a resource that can be used for these purposes. The paper is divided into two parts. First, we analyzed the transcriptomes of 110,814 single RGCs at six ages from embryonic day (E)13, when RGCs are newly postmitotic, to adulthood (postnatal day [P]56; mice are sexually mature by around P40). We previously used these datasets to define 45 mouse RGC types (Tran *et al.*, 2019), and analyze their diversification from a limited repertoire of postmitotic precursors (Shekhar et al., 2022). Here, we surveyed shared and type-specific gene expression changes to identify candidate regulators of RGC development. We identify genes whose temporal regulation overlaps with different phases of RGC development such as differentiation, axonal and dendritic elaboration, synaptic recognition and signaling.

In the second part, we used three models of visual deprivation to assess the effects of light-driven activity on RGC maturation. They were: (1) dark-rearing from birth to adulthood; (2) a well-characterized mutant line (*rd1*) in which visual signals are undetectable shortly after eye-opening (at P14) due to photoreceptor dysfunction and death (Farber et al., 1994; Gibson et al., 2013); and (3) a mutant that lacks bipolar interneurons, which convey photoreceptor input to RGCs (*Vsx2*-SE^-/-^) (Norrie et al., 2019). For each of these three models, we used high-throughput single cell RNA-seq (scRNA-seq) to characterize the transcriptomic diversity of RGCs at P56 and compare their gene expression with that of RGCs from age-matched normally reared mice. We find that all 45 RGC types are found in all models, indicating that visual activity (or lack thereof) does not alter the core transcriptional signatures that specify type-identity. However, visual deprivation attenuates the gene expression differences among RGC types, indicating an impact on transcriptomic maturation or maintenance. We then surveyed the gene groups impacted by visual deprivation and found that they share a significant overlap with the temporally regulated programs underlying normal RGC development. Together, these results indicate that while RGC diversification may be largely governed by vision-independent factors, visual activity plays a role in the final stages of cell type maturation.

## MATERIALS AND METHODS

### Experimental Methods

#### Mice

All animal experiments were approved by the Institutional Animal Care and Use Committees (IACUC) at Harvard University. Mice were maintained in pathogen-free facilities under standard housing conditions with continuous access to food and water. Mouse strains used for both scRNA-seq and histology, were: C57Bl/6J (JAX #000664), *rd1* mice with a mutation in the beta subunit of cGMP phosphodiesterase (*Pde6b*) gene, and *Vsx2*-SE^-/-^ mice lacking bipolar cells (Norrie *et al.*, 2019). The *rd1* mutant had been backcrossed onto a C57Bl/6J background (a kind gift from Prof. Constance Cepko) and the *Vsx2*-SE^-/-^ mice were maintained on a C57Bl/6J background. Embryonic and early post-natal C57Bl/6J mice were acquired either from Jackson Laboratories (JAX) from time-mated female mice or time-mated in-house. For timed-matings, a male was placed with a female overnight and removed the following morning, this being E0.5. The day of birth is denoted P0. For dark-rearing (DR) experiments, animals were kept in light-tight housing in a dark room from the day of birth.

#### Droplet-based single-cell RNA-sequencing of adult RGCs (scRNA-seq)

For each VD condition, RGCs from two separate groups of mice were profiled. Identical protocols were used to isolate RGCs from dark-reared (DR), *rd1* and *Vsx2*-SE^-/-^ mice at P56 and profiled using the same methods described previously for normally reared mice (Shekhar *et al.*, 2022; Tran *et al.*, 2019), with one exception. Retinas from DR mice were dissected in a dark room using a microscope fitted with night-vision binocular goggles (Tactical Series G1, Night Owl), and an external infrared light source, with dissociation and staining steps conducted in LiteSafe tubes (Argos Technologies) to protect from light exposure.

For RGC collection, all solutions were prepared using Ames’ Medium with L-glutamine and sodium bicarbonate, and subsequently oxygenated with 95% O2 / 5% CO2. Retinas were dissected out in their entirety immediately after enucleation and digested in ∼80U of papain at 37°C, followed by gentle manual trituration in L-ovomucoid solution. We used a 40μm cell filter to remove clumps, and following this, the cell suspension was spun down and resuspended in Ames + 4% Bovine Serum Albumin (BSA) solution at a concentration of 10^6^ per 100μl. All spin steps were conducted at 450g for 8 minutes in a refrigerated centrifuge. 0.5μl of 2μg/μl anti-CD90 conjugated to various fluorophores (Thermo Fisher Scientific) was added per 100μl of cells. After a 15 min incubation, the cells were washed with an excess of media, spun down and resuspended in Ames + 4% BSA at 7 x 10^6^ cells per 1 ml concentration. Calcein blue was then added to cells. During fluorescent activated cell sorting (FACS), calcein blue was used to exclude cellular debris, doublets and dead cells, and RGCs were collected based on high CD90 expression.

Following collection, RGCs were spun down and resuspended in PBS + 0.1% non-acetylated BSA at a concentration range of 2000 cells/μl for scRNA-seq processing per manufacturer’s instructions (10X Genomics, Pleasanton, CA). The single-cell libraries for normally reared, *rd1* and DR mice were prepared using the single-cell gene expression 3’ v2 kit on the 10X Chromium platform, while the Vsx2-SE^-/-^ libraries were prepared using the v3 kit. scRNA-seq library processing was done using the manufacturer’s instructions. Libraries were sequenced on the Illumina HiSeq 2500 platforms (paired end: read 1, 26 bases; read 2, 98 bases).

#### Histology and Imaging

Eyes were collected from animals intracardially perfused with 15-50ml of 4% paraformaldehyde (PFA), and post-fixed for an additional 15 min. Dissected eyes and lenses were visually inspected for signs of damage before proceeding further. Healthy eyes were transferred to PBS until retinal dissection, following which retinas were sunk in 30% sucrose, embedded in tissue freezing media and stored at −80°C. Later, retinas were sectioned at 20-25μm in a cryostata and mounted on slides (Tran *et al.*, 2019). Staining solutions were made up in PBS plus 0.3% Triton-X and all incubation steps were carried out in a humidified chamber. Following a 1 hour protein block in 5% Normal Donkey Serum at room temperature, slides were incubated overnight at 4°C with primary antibodies, washed 2 times 5 minutes in PBS, and incubated for 2 hours at room temperature with secondary antibodies conjugated to fluorophores (1:1,000, Jackson Immunological Research), and Hoechst (1:10,000, Life Technologies). Following this incubation, slides were washed again 2 times 5 minutes in PBS and coverslipped with Fluoro-Gel (#17985, Electron Microscopy Sciences).

Antibodies used in this study were guinea pig anti-RBPMS (1:1,000, #1832-RBPMS, PhosphoSolutions), goat anti-VSX2 (1:200, #sc-21690, Santa Cruz Biotechnology), mouse anti-Glutamine synthase (1:1,000, BD Bioscience, # 610517). All images were acquired using an Olympus Fluoview 1000 scanning laser confocal microscope, with a 20x oil immersion objective and 2x optical zoom. Optical slices were taken at 1µm steps. Fiji (Schindelin et al., 2012) was used to pseudocolor each channel and generate a maximum projection from image stacks. Brightness and contrast were adjusted in Adobe Photoshop.

#### Physiology

Mice (C57/Bl6 and *Vsx2*-SE^-/-^) were dark adapted overnight and sacrificed according to protocols approved by the Administrative Panel on Laboratory Animal Care at the University of Washington. After hemisecting each eye, we removed the vitreous mechanically and stored the eyecup in a light-tight container in warm (∼32°C), bicarbonate-buffered Ames Solution (Sigma-Aldrich, St. Louis) equilibrated with 5% CO_2_ / 95% O_2_. All procedures were carried out under infrared light (>900 nm) to keep the retina dark adapted. All experiments were performed in a flat mount preparation of the retina. We placed a piece of isolated retina ganglion cell-side up on a polylysine coated cover slip within a recording chamber. The retina was secured by nylon wires stretched across a platinum ring and perfused with warm (30-34°C) equilibrated Ames solution flowing at 6-8 mL/min. Light from a light-emitting diode (LED; peak output = 470 nm; spot diameter 0.5 mm) was focused on the retina through the microscope condenser.

RGC spike responses were recorded in the cell-attached configuration. RGC inhibitory synaptic inputs were recorded in the voltage-clamp configuration with a holding potential near +10 mV (determined empirically for each cell to eliminate spontaneous inward currents). We used an internal solution containing 105 mM CsCH_3_SO_3_, 10 mM TEA-Cl, 20 mM HEPES, 10 mM EGTA, 5 mM Mg-ATP, 0.5 mM Tris-GTP, and 2 mM QX-314 (pH ∼7.3, ∼280 mOsm); 0.1 mM alexa-488 or alexa-555 was included so that we could image RGC dendrites after recording and identify On and Off cells based on the level of stratification in the inner plexiform layer.

### Computational Methods

#### Normally-reared RGC datasets

Gene Expression Matrices (GEMs) for normally reared (NR) RGCs for E13, E14, E16, P0, P5 and P56 were downloaded from our previous studies (Shekhar *et al.*, 2022; Tran *et al.*, 2019), Entries of raw GEMs reflect the number of unique molecular identifiers (nUMIs) detected per gene per cell, which is a proxy for transcript copy numbers. Raw GEMs were filtered, normalized and log-transformed following standard procedures described before (Shekhar *et al.*, 2022). For each RGC in this dataset, we retained metadata corresponding to cluster/type IDs and also fate associations computed using WOT in (Shekhar *et al.*, 2022), which allowed us to identify putative type-specific precursors at each age. These data can be downloaded from our Github repository: https://github.com/shekharlab/RGC_VD.

#### Force-directed layout embedding of developing RGCs

We visualized the developmental heterogeneity and progression of RGCs on a 2D force-directed layout embedding (FLE) using SPRING (Weinreb et al., 2018) (https://github.com/AllonKleinLab/SPRING\_dev) (Figure 2). To keep the run time manageable, we downsampled our normally reared RGC dataset to 30,000 cells chosen randomly (E13:1661, E14: 4743, E16: 3753, P0: 4968, P5: 4837, P56:10038). The input to SPRING was a matrix of cells by principal component (PC) coordinates computed from the filtered GEMs as follows. To ameliorate within-age batch-effects, we tested RGCs within each age for genes that were globally differentially expressed (fold-difference > 2) within any of the biological replicates. We then computed highly-variable genes (HVGs) in the reduced dataset using a Poisson-Gamma model (Pandey et al., 2018), which resulted in 845 HVGs. The raw GEMs of 845 HVGs by 30,000 cells was once again median normalized and log-transformed. Using PCA, we reduced the dimensionality of this matrix to 41 statistically significant PCs. The 30,000 cells by 41 PCs matrix was supplied to SPRING, which constructed a *k*-nearest neighbor graph (*k*=30) on the data, and used the ForceAtlas2 algorithm (Jacomy et al., 2014) to compute the FLE coordinates over 500 iterations.

**Figure 2.**
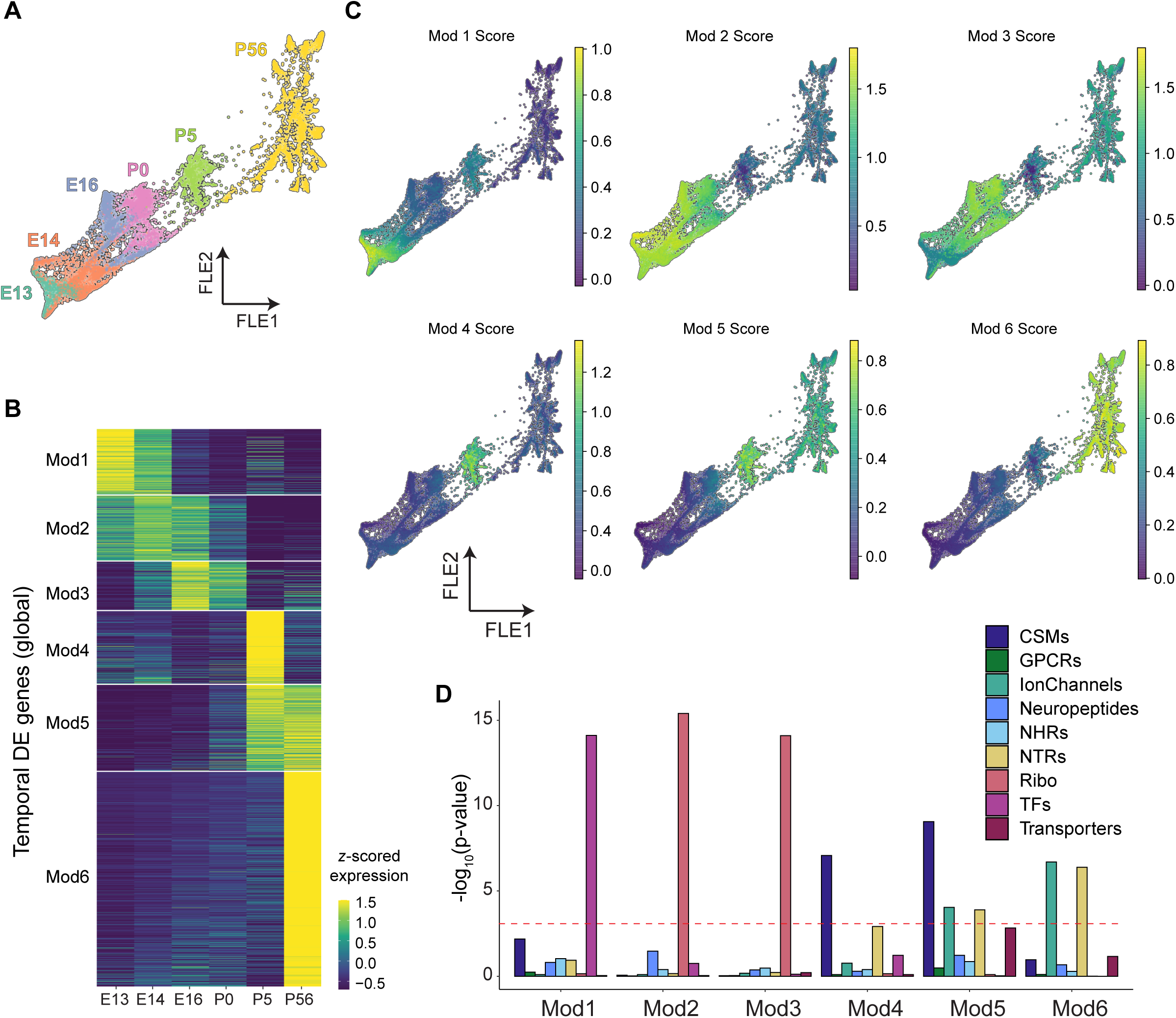
Global gene expression changes during RGC development. A. 2D representation of progressive transcriptomic development of RGCs using a Force-directed layout (FLE) embedding (Weinreb *et al.*, 2018). Individual points represent single RGCs colored by their age. B. Heatmap of temporally regulated genes broadly shared among developing RGCs. Expression values of each gene (row) are averaged across all RGCs at a given age (columns) and z-scored across ages. White horizontal bars separate genes into six modules (Mod1-6) identified by k-means clustering. C. 2D representation as in panel A with individual RGCs colored based on average expression of genes enriched in Mods1-6 (**Table S1**). D. Barplot showing enrichment of cell surface molecules (CSMs), G-protein coupled receptors (GPCRs), ion channels, neuropeptides, nuclear hormone receptors (NHRs), neurotransmitter receptors (NTRs), ribosomal genes (Ribo), transcription factors (TFs) and transporters among Mod1-6. Note that our list of CSMs excludes GPCRs, ion channels, NHRs and NTRs, which are captured in other groups. Each group of bars represents a module, and bar color indicates gene group. Y-axis shows statistical enrichment in −log10(p-value) units. Bonferroni corrected p-values were calculated according to the hypergeometric test (**see Methods**).

The aggregate expression levels for each of the six gene modules were visualized in the 2D FLE as follows. For each cell and module pair, we computed:

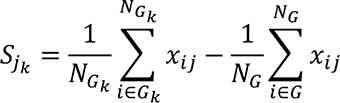

where *S_jk_* is a cell *j*’s score for module *k*, *N_Gk_* is the number of genes in module *k*, *G_k_* is the set of genes in module *k*, *G* is the set of all genes, *N_G_* is the total number of genes, and *x_ij_* is cell *j’s* normalized and log-transformed expression of gene *i.* The first term in the expression reflects the average expression of the module genes in the cell. The second term in the equation subtracts cell *j’s* mean expression across all genes from its module score corrects for baseline differences in library size across cells, a well-known source of variation in scRNA-seq. These scores were used to color cells in the FLE to visualize module activity (Figure 2).

#### Global temporal gene expression changes in developing RGCs

Genes expressed in fewer than 20% of the cells at all the six ages (E13, E14, E16, P0, P5, P56) were discarded. For each remaining gene, the average expression strength *S_g,t_* was computed at each of the six ages as,

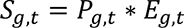

where, *P_g,t_* is the fraction of RGCs at time *t* that express gene *g* (nUMIs > 0) and *E_g,t_* is the log-average expression counts of gene *g* in the expressing cells. We further removed genes that satisfied the criteria,

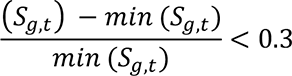

to only consider genes that exhibited > 30% change in expression strength temporally. Next, we randomized the data by shuffling the age labels across RGCs, and used this to compute a “randomized” average expression strength of each gene 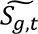. We then ranked genes based on values of the following quantity,

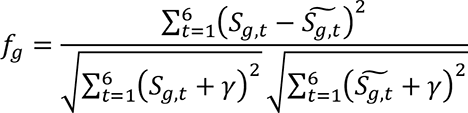

Genes with high values of *f_g_* exhibit greater temporal variability. Here *γ* represents a pseudo-count, chosen to be 0.1 to avoid erroneous inflation of scores for lowly expressed genes. To assess the significance of *f_g_* we computed a null distribution of 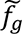 using two different randomizations,

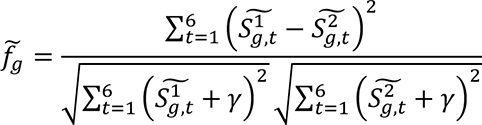

We selected genes that satisfied *f_g_*> *percentile* 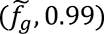. This led to the identification of 1,707 temporally regulated genes. We used k-means clustering to cluster the *S_g,t_* matrix comprised of these 1,707 genes, with the number of groups determined using the gap statistic (Tibshirani et al., 2001). This analysis identified six groups with distinct temporal expression patterns.

#### Gene Ontology (GO) enrichment analysis

We assessed the biological significance of the temporal gene expression modules by performing a Gene Ontology (GO) enrichment analysis. Using the R package topGO (Alexa and Rahnenführer, 2009), each module was evaluated for enrichment of GO terms associated with the three ontology categories: Biological Processes (BP), Molecular Function (MF) and Cellular Component (CC). GO terms with FDR adjusted p-values less than 10^-3^ were identified for each module, and differentially enriched modules were visualized as heatmaps.

#### Enrichment of function gene groups in modules

We assembled lists of mouse transcription factors (TFs), neuropeptides, neurotransmitter receptors (NTRs), and cell surface molecules (CSMs) from the panther database (pantherdb.org). Genes encoding G-protein coupled receptors (GPCRs), ion channels, nuclear hormone receptors (NHRs), and transporters were downloaded from https://www.guidetopharmacology.org/. The list of CSMs were pruned for duplicates by removing genes that were NTRs, GPCRs, NHRs and Ion channels. All genes starting with “Rps” or “Rpl” were tagged as ribosome-associated genes. Each of these gene groups were filtered to only contain genes detected in our dataset. To assess the statistical enrichment of a gene group *g* within a module *m*, we computed the hypergeometric p-value,

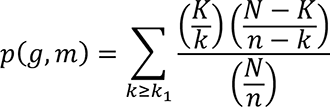

Here *N* is the number of genes in the data, *K* is the number of genes in the group *g, n* is the number of genes in the module *m*, and *k_1_* is the number of genes common between the module *m* and group *g.p*(*g*, *m*) represents the probability that *k_1_* or more genes from the group could be observed in the module purely by random sampling, the null hypothesis. Consequently, low values of *p*(*g*, *m*) or high values of − log *p*(*g*, *m*) are suggestive of significant statistical enrichment.

#### Preprocessing and clustering analysis of single-cell RNA-seq data to recover RGCs

Fastq files corresponding to single-cell RNA-seq libraries from the three VD mice were aligned to the mm10 transcriptomic reference (*M. musculus*) and gene expression matrices (GEMs) were obtained using the Cell Ranger 3.1 pipeline (10X Genomics). GEMs from each of the 12 VD libraries were combined and filtered for cells containing at least 700 detected genes. This resulted in 75,422 cells of which 23,433 were from dark-reared mice, 23,989 were from rd1 mice, and 28,000 were from *Vsx2*-SE^-/-^ mice.

Following standard procedures described previously, the GEMs were normalized and log-transformed, and highly variable genes (HVGs) in the data were identified (Pandey *et al.*, 2018; Shekhar *et al.*, 2022). Within the reduced subspace of HVGs, the data was subjected to dimensionality reduction using PCA and the PCs were batch-corrected across experimental replicated using Harmony (Korsunsky et al., 2019). Using the top 40 PCs, we performed graph clustering (Blondel et al., 2008) and annotated each of the clusters based on their expression patterns of canonical markers for retinal subpopulations described previously (Macosko et al., 2015). The predominant subpopulations included RGCs (82%), amacrine cells (ACs; 12.8%), photoreceptors (4.4%) and non-neuronal cells (0.8%). RGCs were isolated based on the expression of *Slc17a6, Rbpms*, *Thy1*, *Nefl*, *Pou4f1-3*. Overall, we obtained 19,232 RGCs from dark reared mice, 14,864 RGCs from *rd1* mice, and 22,083 RGCs from *Vsx2*-SE^-/-^ mice. RGC yield varied among the three conditions, being 85% for dark rearing, 63.6% for rd1 and 79.3% for Vsx2-SE^-/-^. Following the *in silico* purification of RGCs (Figure S2A), they were subjected to a second round of dimensionality reduction and clustering to define VD clusters (Figures 6, S2).

#### Quantification of transcriptomic separation among RGC types

We used the silhouette score to quantify the degree of separation among RGCs in PC space. Assuming the data has been clustered, let

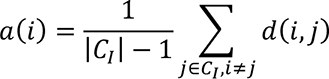

be the mean distance between cell *i* ∈ *C*_*I*_ (cell *i* in cluster *C*_*I*_) and all other cells in the same cluster, where |*C*_*I*_| is the number of cells in cluster *I* and *d*(*i*, *j*) is the distance between cells *i* and *j* in the cluster *I*. We can interpret *a*(*i*) as the average distance of cell *i* is from other members of its cluster. Similarly, we can define the distance of cell *i* to a different cluster *C_j_* as the mean of the distance from cell *i* to all cells in *C_j≠*I*_*. Next, define for each cell *i* ∈ *C*_*I*_

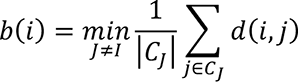

to be the smallest mean distance between cell *i* and the cells of any cluster *C_j≠*I*_*. Finally, the silhouette score for each cell *i* is defined as,

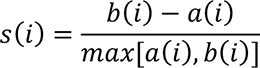

and −1 ≤ *s*(*i*) ≤ 1. *s*(*i*) is a measure of how tightly grouped cell *i* is with other members of the same cluster. The smaller the value of *s*(*i*), the tighter the grouping. Values of *s*(*i*) across the dataset quantify the overall tightness of the clusters. Values near 0 reflect poorly separated, overlapping clusters, while values near 1 indicate highly distinct and well-separated clusters. Negative values indicate that a cell has been assigned to the wrong cluster, as it is similar to cells of a different cluster than to cells of its own cluster.

#### Clustering of VD RGCs and transcriptomic mapping of VD RGCs to control RGC types

RGCs in each VD group were processed separately using the pipeline described above to identify transcriptomically distinct clusters. We then mapped each VD RGC to a normally-reared (NR) RGC type using a classification approach. We used gradient boosted decision trees as implemented in the Python package xgboost to learn transcriptional signatures corresponding to the 45 NR types and used this classifier to assign each VD RGC a NR type label. Three separate classifiers were trained on the NR types, each trained using common HVGs between the atlas and a VD condition as the features. For training, we randomly sampled 70% of the cells in each NR type up to a maximum of 1000 cells. The remaining NR cells were used as “held-out” data for validating the trained classifier to ensure a per-type misclassification rate of less than 5%. Jupyter Notebooks detailing the analysis can be found on https://github.com/shekharlab/RGC_VD.

The trained classifiers were then applied to each VD RGC. To avoid spurious assignments, we only assigned a VD RGC to a type if the classifier voting margin, defined as the proportion of trees accounting for the majority vote, was higher than 12.5%. This is quite stringent as for a multiclass classifier assigning each data point to each of 45 types, a simple majority could be achieved at a voting margin of 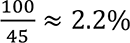. Encouragingly >98% of VD RGCs could be unequivocally assigned to a single RGC type by this criterion. The final mapping between VD clusters and control types were visualized as confusion matrices (Figure S2).

#### Identification of global and type-specific visual-experience mediated DE (vDE) genes

VD-related globally differentially expressed (vDE) genes were evaluated for each visual deprivation condition by VD RGCs as a group with normally reared (NR) P56 RGCs. We used the MAST test (Finak et al., 2015). A gene was considered globally vDE if it exhibited a fold-difference of at least 1.5 and was expressed in at least one condition in >70% all RGCs. We used the same procedure to identify globally DE genes between NR P5 RGCs and NR P56 RGCs. To identify type-specific vDE genes, we repeated the above procedure to each of the 45 RGC type between each of the 3 VD condition and NR - a total of 45*3 = 135 tests. A gene was considered type specific vDE if it was not globally vDE, exhibited a fold-difference of at least 1.5 in a type across conditions, and was expressed in at least 30% of cells of that type in either condition.

## RESULTS

Mouse RGCs are the first-born neuronal class in the retina, with >95% of RGCs arising between embryonic days (E) 12 and 17 (Drager, 1985; Farah and Easter, 2005; Marcucci et al., 2019; Voinescu et al., 2009). RGC axons begin reaching retinorecipient targets by E15 and are refined in an activity-dependent manner postnatally as noted above (Godement et al., 1984; Osterhout et al., 2011). Dendritic development, however, only commences during early postnatal life, when RGCs receive synapses in the inner plexiform layer (IPL) from amacrine cells by P4 and from bipolar cells soon thereafter (Kim et al., 2010; Lefebvre et al., 2015; Sernagor et al., 2001). Beginning around E16, RGCs exhibit spontaneous and synchronized activity that propagates in a wave-like fashion (Feller and Kerschensteiner, 2020). Initially, activity is light-independent but starting around P10, light penetrating the eyelids generates responses in the photoreceptors that are transmitted to RGCs (Tiriac *et al.*, 2018). Image-forming vision begins after P14, when the eyes open (Hooks and Chen, 2007) (Figure 1B). By the time of eye-opening, RGCs exhibit diverse type-specific axonal and dendritic arborization patterns (Kim *et al.*, 2010), and feature-selective responses can be recorded *ex vivo* (Tiriac et al., 2022), indicating that RGC types and their basic circuitry are established without image-forming visual experience.

### RGC diversification

To study RGC diversification we recently used scRNA-seq to profile RGCs at E13, E14, E16, P0, P5, and P56 (Shekhar *et al.*, 2022; Tran *et al.*, 2019). We briefly summarize the main conclusions of that study before proceeding to describe changes in gene expression during this period.

First, the number of molecularly distinct groups of RGCs as well as their transcriptomic distinctiveness increases with age. Groups of RGCs at the earliest stages studied exhibit graded gene expression differences spanning a transcriptomic continuum, but they become increasingly discrete as development proceeds. Second, using a computational method called Waddington optimal transport (WOT; (Schiebinger et al., 2019); described below), we found that RGC types are not specified at the progenitor level but rather that multipotentiality persists in postmitotic precursor RGCs. Third, these precursor RGCs become gradually and asynchronously restricted to specific types with developmental age. Finally, diversification may in many cases occur in two steps, with precursors initially committing to subclasses, each defined by the selective expression of TFs in the adult. Subsequently, precursors within a subclass become restricted to single types by a process we named “fate-decoupling” (Shekhar *et al.*, 2022).

### Gene expression changes as RGCs develop

To analyze gene expression patterns during RGC development, we first combined all single cell transcriptomes at each age to identify changes broadly shared among types. We visualized the developmental progression at single-RGC resolution using a force-directed layout embedding (FLE; Figure 2A). FLE arranges cells on a 2D map based on mutually attractive/repulsive “forces” that depend on transcriptional similarity (Fruchterman and Reingold, 1991; Jacomy *et al.*, 2014). We then identified 1,707 global temporally-regulated DE genes and used *k*-means clustering to group them into modules with distinct temporal dynamics, choosing k=6 based on the gap-statistic method (see **Methods**; Figures 2B; **Table S1**). Figure 2C shows the FLE plot with each cell colored based on the average expression levels of genes in each of the six modules, verifying that the expression of these modules varied systematically with age and were broadly shared among RGCs.

We analyzed the modules (Mod1-6) in two ways. First, we examined the enrichment of nine functional groups of genes: cell surface molecules (CSMs), G-protein coupled receptors (GPCRs), ion channels, neuropeptides, nuclear hormone receptors (NHRs), neurotransmitter receptors (NTRs), ribosomal genes (Ribo), transcription factors (TFs) and transporters (Figure 2D). Second, we used conventional gene ontology (GO) enrichment analysis to highlight distinct biological processes, molecular functional units and cellular components that were enriched in each of these modules (Figure 3)(The Gene Ontology, 2019).

**Figure 3.**
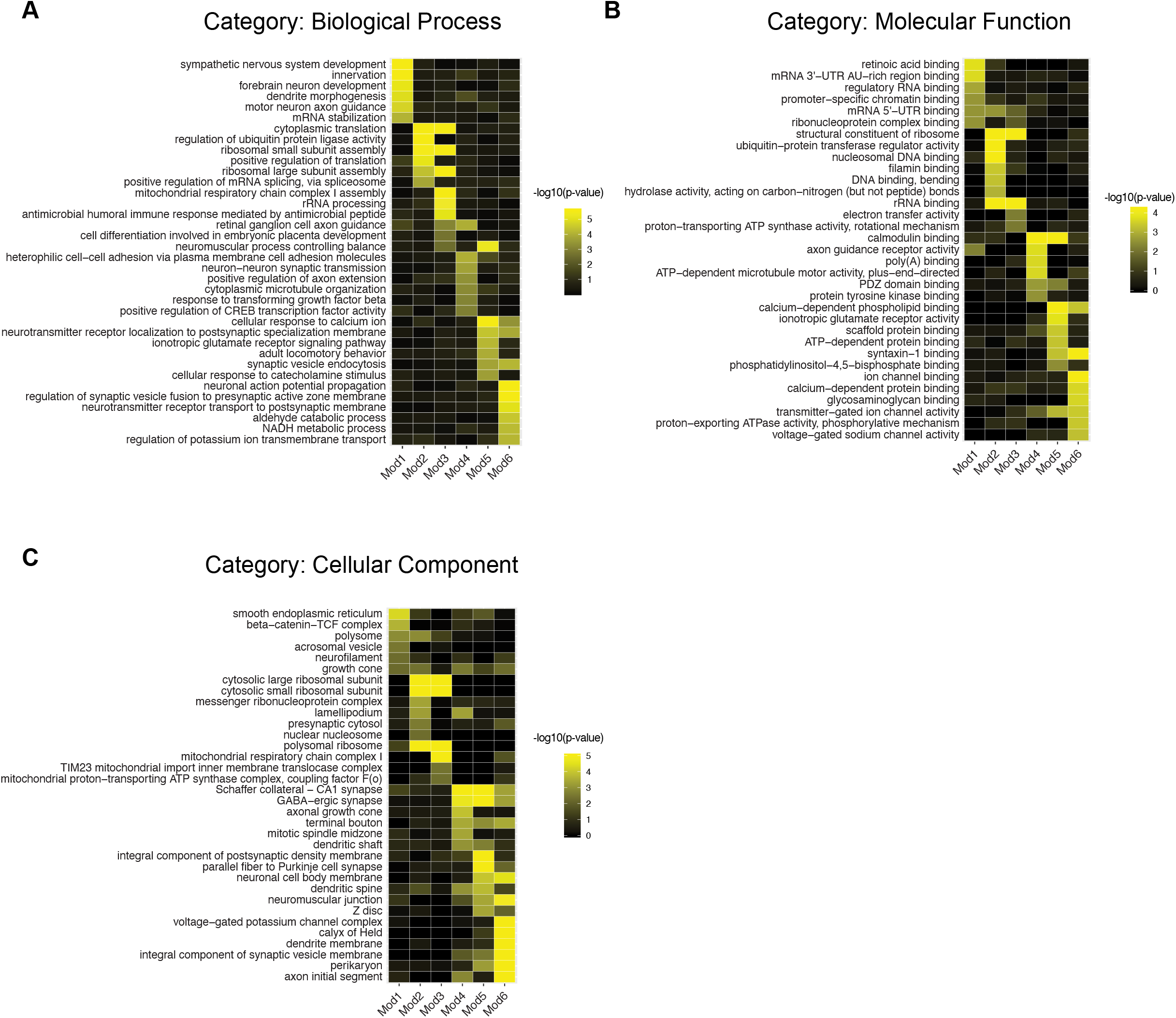
Gene Ontology (GO) analysis of temporally regulated gene modules Mod1-6. A. Examples of significantly enriched Gene Ontology (GO) category “Biological Process (BP)” terms (rows) in Mod1-6 (columns). Colors correspond to enrichment corrected p-values (-log10 units). GO analaysis was performed using the R package topGO (Alexa and Rahnenführer, 2009). B. Same as A, for GO category “Molecular Function (MF)”. C. Same as A, for GO category “Cellular Component (CC)”.

Mod1, consisting of 202 genes, was most active at E13. It included genes associated with retinoic acid signaling (*Crabp1, Crabp2, Nr2f2*) as well as axon guidance (*Robo1, Nrp1, Sema3a, Slit1*), coinciding with the initial period of axon growth (Zhang et al., 2017) (Figures 3A,B). Mod1 was also the only module significantly enriched for TFs (Figure 2D). TFs included several previously shown to influence RGC differentiation, such as SoxC class TFs *Sox11/12* (Jiang et al., 2013; Kuwajima et al., 2017), *Onecut2/3* (Sapkota et al., 2014), *Nhlh2*, *Ebf3* and *Irx3,5* (Jin et al., 2010; Lu et al., 2020; Lyu and Mu, 2021). We also found many TFs that have not, to our knowledge, been previously described in this context, including *Baz2b, Hmx1, Tbx2* and *Zeb2*.

Expression of genes in Mod2 and Mod3 was highest at later prenatal ages - E14 for Mod2 and E16 for Mod3. Genes enriched in these modules included those associated with ribosome biogenesis and assembly (Rps-genes and Rpl-genes), translation, and mitochondrial function (*Ndufa1-3, Ndufb2,4, Ndufv3*), all consistent with requirements for neuronal growth and maturation during this period (Figures 2D and 3C).

Mod4-Mod6 were most active at the postnatal ages: P5 for Mod4, P56 for Mod6, and both ages for Mod5. Mod4 and Mod5 contained many genes encoding cell surface molecules (Figure 2D). Among them were genes implicated in formation of retinal neural circuits, including members of the three superfamilies most prominently implicated in synaptic specificity: the cadherin (e.g., *Cdh4, Cdh11, Pcdh17/19, Pcdha2, Pcdhga9*), immunoglobulin (e.g., *Dscam, Ncam2*, and *Nrcam*) and leucine-rich repeat super<u>-f</u>amiles (*Lrrn3, Lrrtm2, Lrrc4c*) as well as teneurins (*Tenm1, Tenm2, Tenm4*), which are counter-receptors for leucine-rich repeat proteins (Sanes and Zipursky, 2020). Mod5 was enriched for transporters (e.g. *Slc6a1, Slc6a11, Slc24a3, Atp1a1, Atp1a3*). All three postnatal modules were enriched for genes required for synaptic transmission, such as GABA receptors (*Gabra3, Gabrbr3, Gabbr2*), ionotropic glutamate receptors (*Gria1-4* and *Grin1/2b*), and synaptotagmins (*Syt1,2,6*). Mod5 and Mod6 were especially enriched for genes encoding ion channels including many associated with action potential propagation (*Scn1a, Scn1b, Cacnb4*, *Kcna1, Kcnc2, Kcnip4, Kcnab2;* **see** Figure 3B,C).

Taken together, these data provide a comprehensive catalogue of molecular changes associated with RGC maturation, including genes implicated in neuronal differentiation and growth, axon guidance and synaptogenesis, and acquisition of electrical and synaptic capabilities.

### Type-specific gene expression changes during RGC maturation

We next leveraged the single-cell resolution of our dataset to identify genes selectively expressed in small subsets of RGC precursors. Such genes may contain factors that instruct specific fates (fate determinants), or RGC type-specific properties. To this end, we identified genes that were specific to transcriptomic clusters at each age as defined in our previous studies: they were expressed in at least 30% of the cells in fewer than 10% of the clusters and in no more than 5% of the cells in any remaining cluster (Shekhar *et al.*, 2022; Tran *et al.*, 2019). The number of specific genes increased steadily with age, from 2 at E13 (*Lect1* and *Pou4f3*) to 200 at P56 (**Table S2**). The increase was striking even when taking the increasing number of clusters into account (10 at E13 and 45 at P56; Figure 4A; (Shekhar *et al.*, 2022). The gene categories introduced in Figure 2D accounted for 45-55% of genes at each age. The two most prominent categories were TFs and CSMs, accounting for 12-50% and 8-18% of all genes, respectively (Figure 4B). All the other categories were represented at lower proportions. There was substantial turnover of specific genes with age: only ∼23% of specific genes at E14 and E16 and ∼50% of specific genes at P0 and P5 remained specific at P56, reflecting the dramatic transcriptomic changes that occur during RGC diversification and maturation.

**Figure 4.**
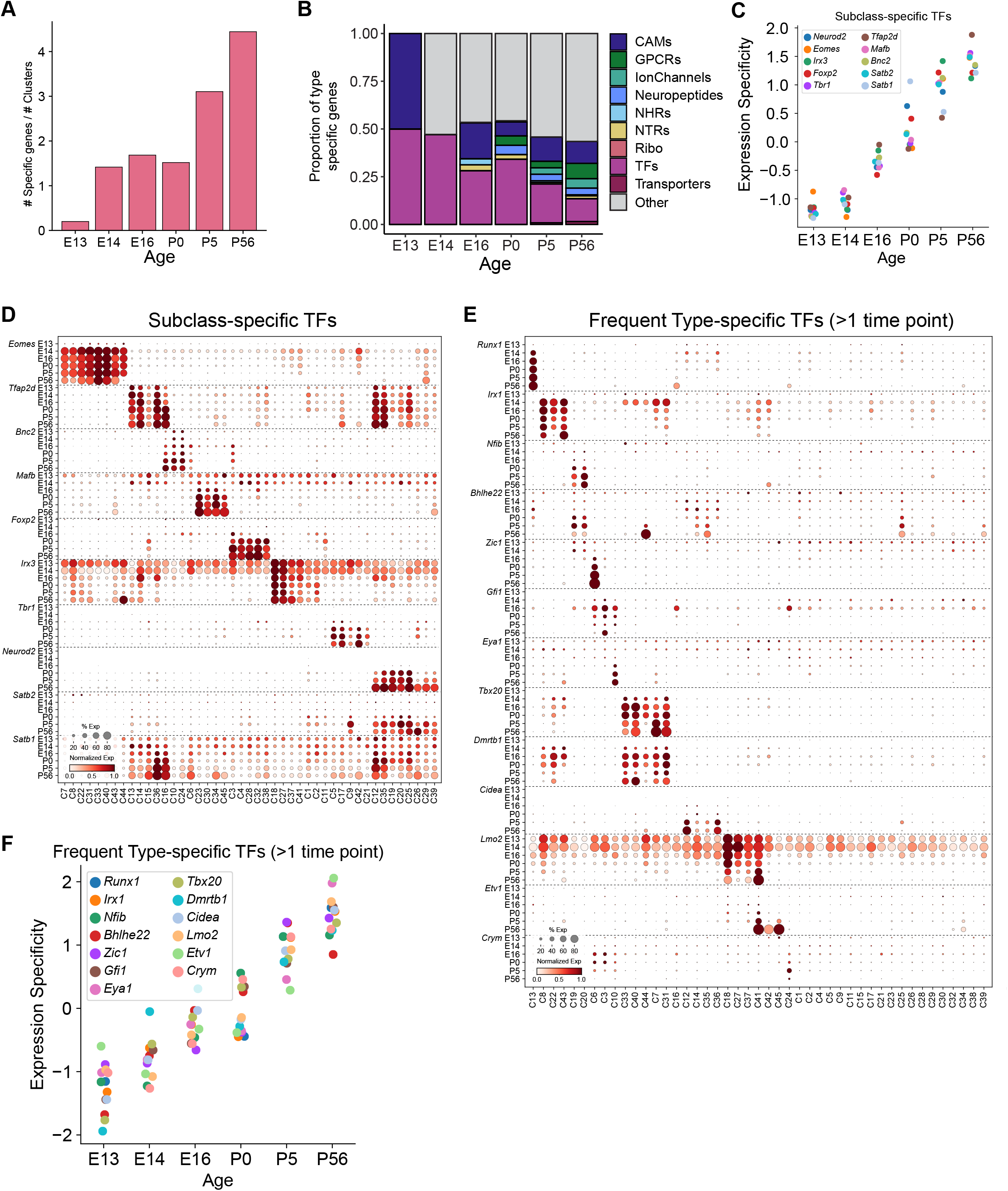
Subclass- and type-specific gene expression changes during RGC development. A. The number of specific genes per cluster increases with age. Y-axis plots the number of type-specific genes divided by the number of clusters at each age, as defined in Shekhar et al., 2022 (10 clusters at E13, 12 clusters at E14, 19 clusters at E16, 27 clusters at P0, 38 clusters at P5, and 45 clusters at P56). B. Relative proportion of each of the 10 gene groups in Figure 2D among the specific genes in panel A. Abbreviations as in Figure 2D. Note that E13 contains only two specific genes: one CSM and one TF. C. Expression of eight RGC subclass-specific TFs (as in Shekhar et al., 2022) becomes increasingly specific with age. Expression specificity is defined as the z-scored dispersion of expression levels across cell types. At ages earlier than P56, putative precursors were inferred using WOT (see **Methods**). D. Dot plot showing expression patterns of subclass-specific TFs with age among putative type-specific precursors. Each row displays the expression levels of a TF at a particular age among type-specific precursors (columns) identified using WOT. The size of a dot corresponds to the fraction of cells with non-zero transcripts, and color indicates normalized expression levels. Row blocks corresponding to different TF are demarcated by dotted horizontal lines. In addition to the eight subclass-specific TFs in C, we also plot the expression patterns two RGC selective TFs, *Satb1* and *Satb2* (Dhande et al., 2019; Peng *et al.*, 2017). E. Same as D, showing the expression of TFs identified as type-specific among WOT-inferred precursors at least two ages via DE analysis. F. Same as C, showing increasing specificity of type-specific TFs plotted in E.

To visualize the temporal evolution of these genes as RGC diversification progressed, we linked cell types across time with Waddington optimal transport (WOT)(Schiebinger *et al.*, 2019). Briefly, WOT uses transcriptomic similarity as a proxy to directly compute fate associations at the level of individual cells without requiring clustering as a prior step, identifying putative precursors of each of the 45 adult RGC types at each of the early time points. This, in turn, enables us to visualize gene expression changes along the inferred developmental history of each type (see (Shekhar *et al.*, 2022) for further discussion and validation).

We used this framework to examine three sets of genes. First, we queried expression of a set of TFs that are expressed in transcriptomically proximate types of adult RGCs that we nominated as subclasses defined by shared-fate association (Shekhar *et al.*, 2022). Many of these transcription factors have been noted in previous analyses as selectively expressed among RGC types (Kiyama et al., 2019; Liu *et al.*, 2018; Mao et al., 2020; Rheaume *et al.*, 2018; Rousso et al., 2016; Tran *et al.*, 2019). Analysis of their expression revealed a gradual increase in expression specificity (Figure 4C-D). This coincides with gradual specification of RGC subclasses observed by Shekhar et al. (2022), with the *Eomes* and *Neurod2* subclasses emerging earliest and latest, respectively. For the few that have been studied functionally, expression patterns were consistent with their roles (see **Discussion**).

Second, we sought TFs that were selectively expressed in just 1-3 inferred types at one or more of the six ages, reasoning that they might include type-specific fate determinants. TFs in this category included *Zic1* in type C6 (nomenclature of (Tran *et al.*, 2019)), *Gfi1* in C3, *Eya1* in C10, *Runx1* in C13, *Msc* in C31 and *Esrrb* in C41 (Figures 4E, S1A). Several of these types have been characterized morphologically and/or physiologically (see **Discussion**) but roles of the TFs remain to be explored. The majority of the selectively expressed TFs exhibited increasingly restricted expression with age, consistent with the overall transcriptomic divergence of RGCs (Figure 4F). However, this trend was not universal: for example, *Esrrg* and *Fgf1* became less specific with age (Figure S1C).

Third, we analyzed expression of recognition molecules that we and others have shown to play roles in synaptic choices of RGCs (Figure S1B) (Duan *et al.*, 2014; Duan et al., 2018; Krishnaswamy *et al.*, 2015; Liu *et al.*, 2018; Liu and Sanes, 2017; Matsuoka et al., 2011; Osterhout *et al.*, 2011; Osterhout et al., 2015; Peng *et al.*, 2017; Yamagata and Sanes, 2018). Most of the genes queried become specific only during postnatal ages (Figures S1D), consistent with the known timing of dendritic elaboration and synaptogenesis, but nonetheless exhibit variability in timing. These data provide a rich resource for identifying candidate molecules that may, likely in combination, regulate selective aspects of RGC type identity.

### Visual deprivation models

To assess the effects of visual experience on RGC maturation we analyzed three groups of adult mice in which in which RGCs had been visually deprived (VD) postnatally. The first model, dark-rearing (DR) from birth to analysis in adulthood (P56), deprives the retina of all postnatal visual input. Standard histology, including immunostaining with antibodies to the RGC-specific marker Rbpms showed that dark-rearing had no obvious effect on retinal structure, and that RGCs were normal in number and position (Figure 5A).

**Figure 5.**
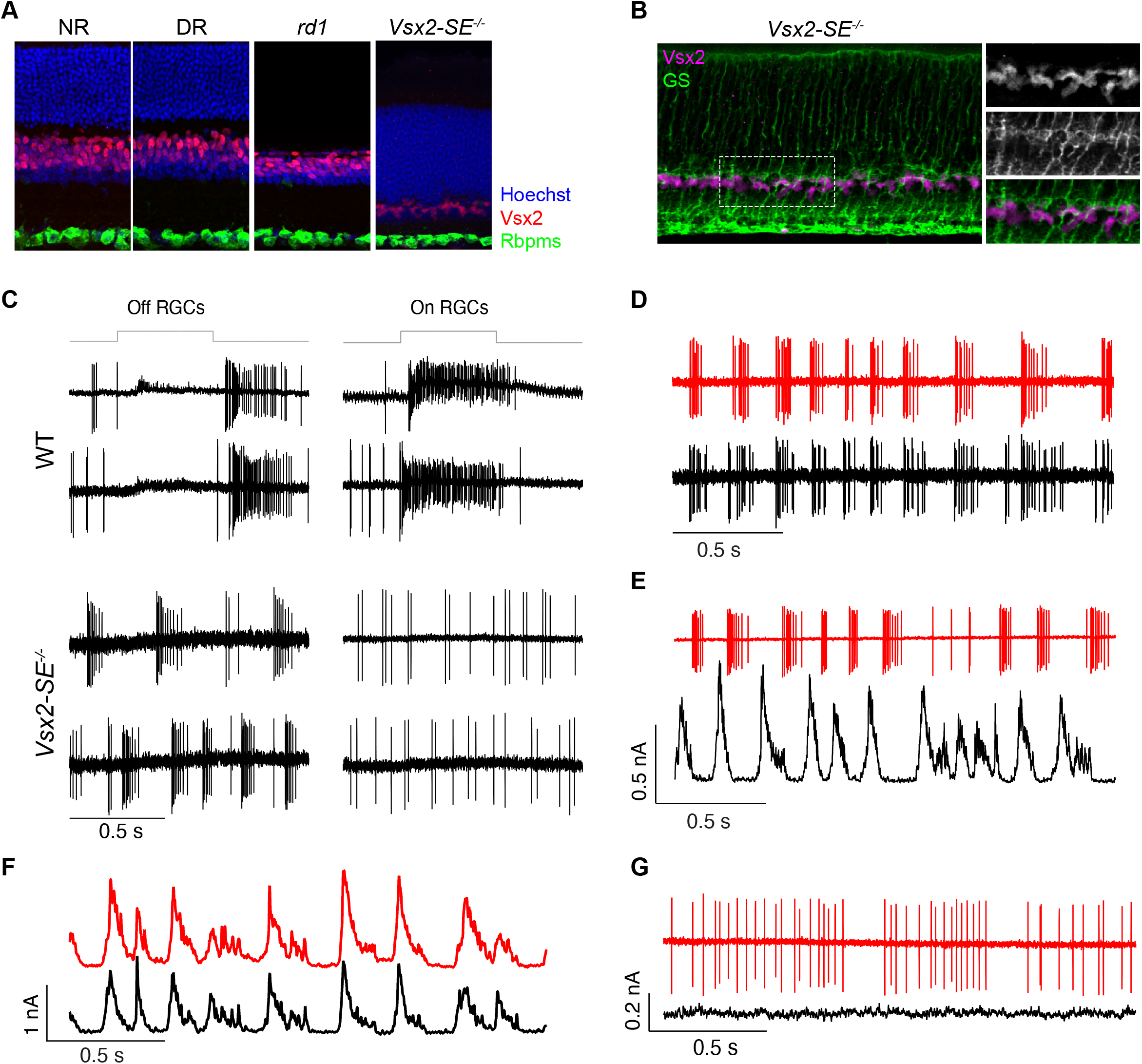
Three visual deprivation models. A. Sections from adult (P56) retinas of normally reared (NR), dark-reared, *rd1* and *Vsx2*-SE^-/-^ mice stained for Vsx2 (a pan-BC and Müller glia marker), Rbpms (pan-RGC marker), and Hoechst (nuclear marker). B. Section from *Vsx2-*SE^-/-^ retina co-stained for glutamine synthetase (GS), a marker of Müller glia, and Vsx2. All Vsx2 immunoreactivity in the mutant retina is associated with Müller glia. C. *Vsx2-*SE^-/-^ RGCs fail to respond to light. Cell-attached recordings of responses to a light step from four wild-type (WT) RGCs (two OFF-sustained and two ON-sustained) and lack of response from four *Vsx2-*SE^-/*-*^ RGCs. Light step for WT RGCs was adjusted to 10 R*/rod/s. Light step for *Vsx2-*SE^-/-^ RGCs was to 1000 R*/rod/s. Note that the pattern of spontaneous activity is very different between OFF and ON *Vsx2*-SE^-/-^ RGCs. R*/rod/s is the photoisomerization rate. D. Strong synchrony between two nearby *Vsx2-*SE^-/-^ OFF RGCs. Simultaneously-recorded OFF RGCs produce highly correlated spontaneous bursts of activity. Colors indicate two different RGCs. E. Increase in inhibitory input produces gaps in spontaneous firing. Panels show simultaneous recordings of spikes in one OFF RGC and inhibitory input in another nearly OFF RGC. Increases in inhibitory input are correlated with a decrease in spontaneous firing in the neighboring cell. F. Inhibitory input to OFF RGCs is strongly correlated. Simultaneous recordings of spontaneous inhibitory input to two nearby OFF RGCs. The correlation coefficient for this pair was 0.8. G. Blocking of GABA (gabazine and TPMPA) and glycine (strychnine) receptors eliminates synchrony and patterned spontaneous activity.

The second model is the well-characterized and widely used *rd1* line, which carries a nonsense mutation in the gene encoding the rod-specific cGMP phosphodiesterase 6-beta subunit (*Pde6b*). Loss of *Pde6b* results in dysfunction and death of rod photoreceptors, which is followed, for reasons that remain largely unknown, by loss of cone photoreceptors (Farber *et al.*, 1994; Keeler, 1924; Punzo and Cepko, 2007). Although rods are not completely lost until one month of age and cones later still, their function is disrupted earlier, and visual signals are undetectable by P21 (Gibson *et al.*, 2013). Since eye-opening does not occur until P14, RGCs in *rd1* mice experience conventional visual input only for a brief period. The outer nuclear layer, which contains rods and cones, was nearly absent from *rd1* retina by P56, but the number of RGCs in the ganglion cell layer was not detectably affected (Figure 5A).

The third model is a mutant (*Vsx2*-SE^-/-^) lacking the bipolar interneurons that convey signals from rod and cone photoreceptors to RGCs ((Gamlin et al., 2020; Norrie *et al.*, 2019); see Figure 1A). *Vsx2* is expressed in retinal progenitor cells and its expression is maintained in differentiated bipolar neurons and Müller glia. It is required for early progenitor divisions and also for formation of bipolar cells (Burmeister et al., 1996; Liu et al., 1994). An enhancer essential for *Vsx2* expression in bipolar cells is deleted in this line, resulting in failure of bipolar cells to form. As the remainder of the *Vsx2* gene is intact, other retinal cell classes form normally. Thus, RGCs receive no visual input in *Vsx2*-SE^-/-^ mice, although it is likely that ipRGCs, which express the photosensitive pigment melanopsin, retain visual responsiveness. As expected, the inner nuclear layer was thin is this mutant, but there was no significant effect on the thickness of the outer nuclear layer, which contains photoreceptors, or the ganglion cell layer in which RGCs reside (Figure 5A). We noted some residual staining with anti-Vsx2 but determined that this reflected the retention of Müller glial cells (glutamine synthetase-positive), which express Vsx2 and are unaffected by deletion of the bipolar-specific enhancer (Figure 5B).

RGCs in the *rd1* line have been characterized previously (Choi et al., 2014; Goo et al., 2015; Stasheff, 2008), but those in the *Vsx2*-SE^-/-^ line have not. We therefore recorded from RGCs in isolated *Vsx2*-SE^-/-^ retinas. RGCs were labeled by inclusion of a fluorescent dye in the recording pipette. We targeted cells with the largest somata, which in wild-type retinas are ON-sustained, OFF-sustained and OFF-transient RGCs. ON or OFF RGCs were identified based on confocal imaging following recording; dendrites of likely ON cells (n=5) arborized near the ganglion cell layer, while dendrites of likely OFF cells (n=10) arborized near the inner nuclear layer.

As expected, none of the recorded RGCs generated measurable changes in firing rate in response to light steps; identical steps elicited large responses in WT RGCs (Figure 5C). *Vsx2*-SE^-/-^ OFF RGCs generated spontaneous rhythmic activity consisting of high-frequency bursts of spikes separated by periods of silence (Figure 5C). Firing rates during the bursts often exceeded 100 Hz. ON RGCs lacked this rhythmic activity, instead generating occasional spontaneous spikes that were not organized into bursts. Dual recordings demonstrated that spontaneous activity was strongly correlated between nearby OFF RGCs (Figure 5D). Similar spontaneous activity in *rd1* mice appears to originate in AII amacrine cells, which provide direct inhibitory input to OFF but not ON RGCs in wild-type mouse retina. Consistent with this mechanism, inhibitory input to an OFF RGC coincided with pauses in firing in a nearby OFF RGC (Figure 5E) and inhibitory input to nearby RGCs was very strongly synchronized (Figure 5F; peak correlation in three pairs was 0.7, 0.8 and 0.9, compared to 0.2-0.3 for pairs of WT RGCs). Moreover, pharmacological blockade of inhibitory synaptic transmission abolished the rhythmicity of activity in OFF RGCs (Figure 5G). RGCs did not show evidence for direct synaptic interactions: depolarizing one cell in a paired recording did not elicit a measurable response in the other (data not shown). These observations support a picture in which synchronized activity in the AII amacrine network produces strongly synchronized inhibitory input to OFF RGCs and produces coordinated pauses in their spontaneous firing.

### Effects of visual deprivation on RGC type identity

To study the influence of visual input on RGC type identity, we obtained scRNA-seq profiles of 19,232 RGCs from DR mice, 14,864 RGCs from *rd1* mice, and 22,083 RGCs from *Vsx2*-SE^-/-^ mice, all at P56 (**Methods**; Figure S2A). We separately clustered each dataset in an unsupervised fashion to identify molecularly distinct RGC clusters (Figure S2B-E). We then used a classification framework (Chen and Guestrin, 2016; Tran *et al.*, 2019) to map each VD RGC to P56 NR RGC types (Chen and Guestrin, 2016; Tran *et al.*, 2019) (**Methods**; Figures 6A-D and S2F-H). This framework employs an ensemble of gradient boosted decision trees, with each decision tree assigning any given VD RGC to one of the 45 adult types in NR mice. A VD RGC was assigned to the NR type that received the majority vote. Cells were considered unequivocally mapped if their voting margin was ≥12.5%, or 5-fold higher than 2.2% (1/45), the margin of a classifier that votes randomly. All 45 types were recovered in all three models, and mapping was highly specific in that 98.5% of the 56,179 VD RGCs mapped to a single type based on this criterion (Figure 6E). Indeed, the average voting margin for a VD RGC was >90% for all three conditions, or 40-fold higher than chance, and the distribution was similar among conditions. Moreover, TFs and adhesion molecules that were type-specific in NR retina retained their type-specificity in all three VD models (Figure S3).

**Figure 6.**
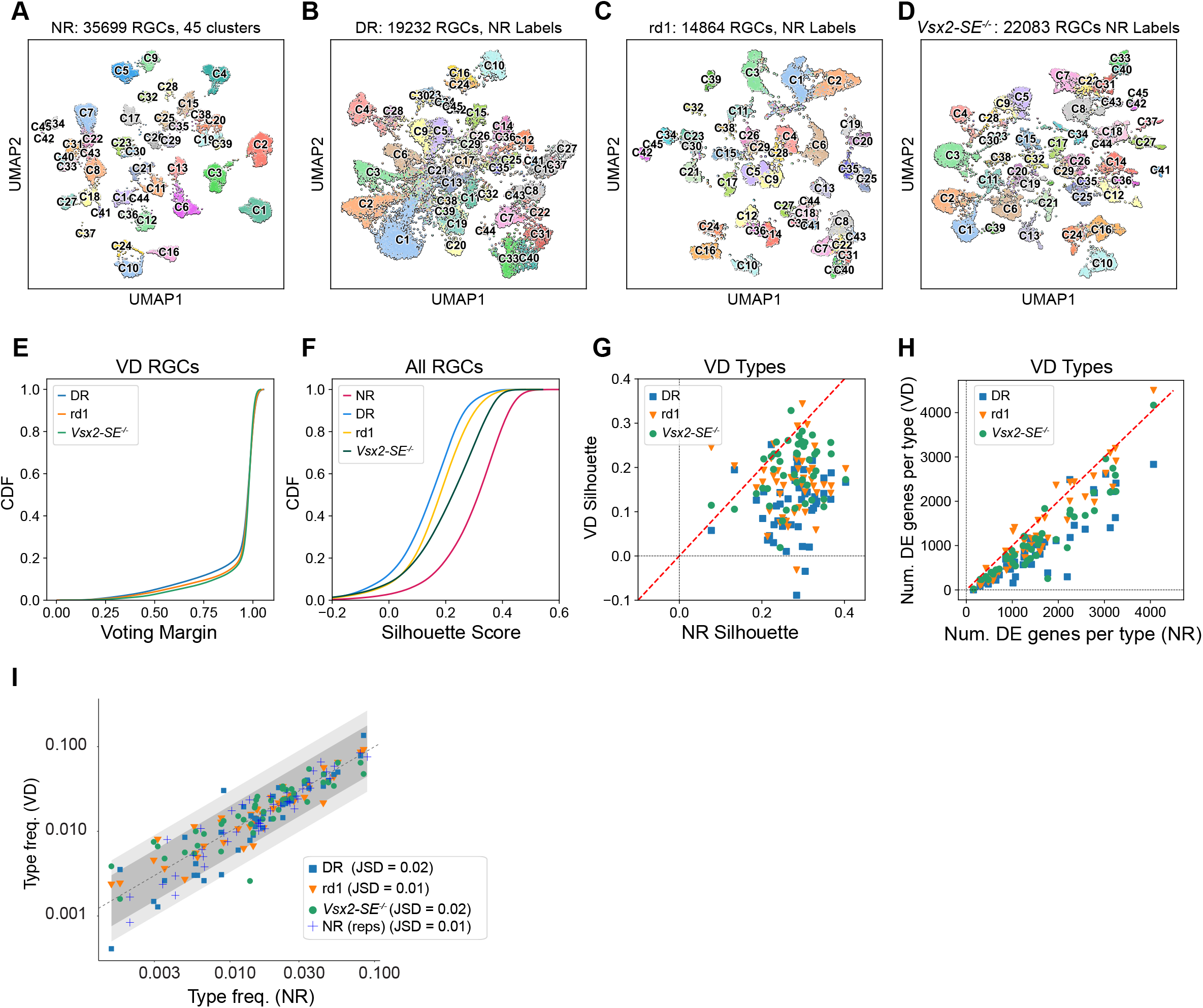
Transcriptomic classification of RGCs from visually deprived mice. A. 2D visualization of the transcriptomic diversity in normally reared (NR) P56 RGCs using Uniform Manifold Approximation (UMAP; (Becht et al., 2019)). Individual cells (points) represent single RGCs, and are colored by type identity as in Tran et al. (2019). B. UMAP visualization DR RGC transcriptomes profiled in this study. Individual RGCs are colored based on NR type-identity as determined using a supervised classification framework (**Methods**). C. Same as B, for *rd1* RGCs. D. Same as B, for *Vsx2-*SE^-/-^ RGCs. E. Cumulative distribution functions (CDFs) showing voting margins for VD RGCs by condition (colors). The voting margin is defined as the fraction of decision trees casting the majority vote. A VD RGC is assigned to the type receiving the majority vote as long as the margin is ≥5X greater than chance, corresponding to a margin of 0.022 (see **Methods**). F. CDFs for silhouette coefficients for NR, DR, *rd1* and *Vsx2-*SE^-/-^ RGCs (colors). Details of calculating the silhouette coefficients are described in the **Methods**. CDFs for VD RGCs were significantly different from the CDF for NR RGCs based on the two-sample Kolmogorov-Smirnov test (p-values are 10^-106^, 10^-88^, and 10^-50^ for DR vs. NR, *rd1* vs. NR, and *Vsx2*^-/-^-SE vs. NR comparison, respectively). G. Comparison of the average silhouette coefficient for each of the 45 RGC types under NR (x-axis) and VD conditions (y-axis). Each point corresponds to a type (45*3=135 total points), and colors and symbols (legend) indicate VD condition. H. Scatter plot showing that there are fewer DE genes per type among VD RGCs (y-axis) than normally reared RGCs (x-axis). Colors and symbols as in G. I. Scatter plot comparing relative frequencies of RGC types in NR (x-axis) vs. VD (y-axis). Colors and symbols of VD as in G. Blue crosses represent frequencies observed in replicates of NR RGCs. Dashed line shows y=x. Shaded ribbons are used to represent a frequency-fold change difference of 2 (dark gray) and 3 (light gray), respectively. JSD - Jensen Shannon Divergence, a measure of distance between the frequency distributions (0 – identical distributions, 1 – maximally disparate distributions). Difference between VD and control are in most cases comparable to those observed between NR replicates.

Although nearly all VD RGCs could be assigned to NR types, two observations led us to examine the influence of visual deprivation on the specification of RGC types: First, VD RGCs were less transcriptomically separated in UMAP projections than their NR counterparts (Figure 6A-D). Second, when we assessed the correspondence of VD clusters to NR RGC types there were several cases in which not all RGCs within a single VD cluster mapped to the same NR type; instead, RGCs mapping to 2 or 3 different types co-clustered in the VD dataset (Figure S2F-H). To evaluate decreased transcriptomic separation as an explanation for multimapping, we calculated for each RGC in each condition, the silhouette score, a measure of the tightness of clustering in principal component space (Rousseeuw, 1987). The silhouette score for an RGC is a measure of how similar it is transcriptomically to other RGCs of the same type compared to RGCs of other types (**Methods**). VD RGCs exhibited consistently lower silhouette scores than their NR counterparts (Figure 6F) and nearly all types exhibited lower average silhouette scores compared to their NR counterparts (Figure 6G). Subsampling analyses verified that these differences were not driven by the larger sample size of NR RGCs. Moreover, RGCs in “multimapped” clusters generally belonged to the most transcriptomically similar types in the NR retina (Tran *et al.*, 2019). Consistent with the failure to fully acquire or maintain type-specific distinctions, >90% of RGC types in the VD conditions exhibited fewer DE genes than their NR counterparts (Figure 6H). Taken together, these results indicate that RGCs acquire their type identity in a vision-independent manner but require visual input for complete transcriptomic maturation or maintenance.

We also compared the relative frequencies of each RGC type in NR and VD models. 40/45 types exhibited less than 2-fold change in relative frequency compared to the atlas across the full range of observed frequencies (0.1% to 8%). Such changes were comparable to those observed between normally reared P56 biological replicates (Figure 6I), suggesting that they likely represent sampling variation rather than true biological changes. Further the frequency distributions of types between VD and NR RGCs were very similar, as quantified by near zero values of the Jensen-Shannon divergence (JSD), a measure of divergence between two frequency distributions (Bishop and Nasrabadi, 2006). Although we cannot rule out the possibility that larger samples and more replicates would reveal modest changes in type frequency, we conclude that VD has no significant differential effect on the generation or maintenance of specific RGC types.

### Effects of visual deprivation on gene expression

Finally, we compared RGCs from each VD condition to NR RGCs to identify visual-experience dependent DE (vDE) genes (**Methods**). We began by identifying transcriptomic alterations that were broadly shared among RGCs (global vDE) (Figure S2I). We found a total of 477 genes that exhibited a >1.5-fold change between NR and at least one VD condition (MAST DE test; adjusted p-value < 10^-4^) and were detected in at least 70% of RGCs in either condition. At the bulk level, the transcriptomic profiles of RGCs from all three VD conditions were more similar to NR RGCs at P56 and to each other than they were to NR RGCs at P5 (Figure 7A). The number of global DE genes between P5 and P56 control RGCs, defined using identical metrics, is several-fold larger than the number of global vDE genes at P56 between NR RGCs and any of the VD conditions. This difference is easily appreciated from the elliptical rather than circular profiles when gene expression changes between P5 NR and P56 NR RGCs are compared to those between P56 NR and P56 VD RGCs (Figures 7A-C). As an example, 217 genes are down-regulated and 759 up-regulated in NR RGCs between P5 and P56, whereas only 23 (11% as many) are down-regulated and 161 (21% as many) up-regulated in dark-reared compared to NR mice at P56 (Figure 7B). This difference is insensitive to the choice of DE threshold in the range 1.2-fold to 1.8-fold, and very few genes exhibit higher fold changes in VD. Based on these results, we conclude that the majority of gene expression changes that occur during the maturational period between P5 and P56 do not rely on visual experience-driven activity.

**Figure 7.**
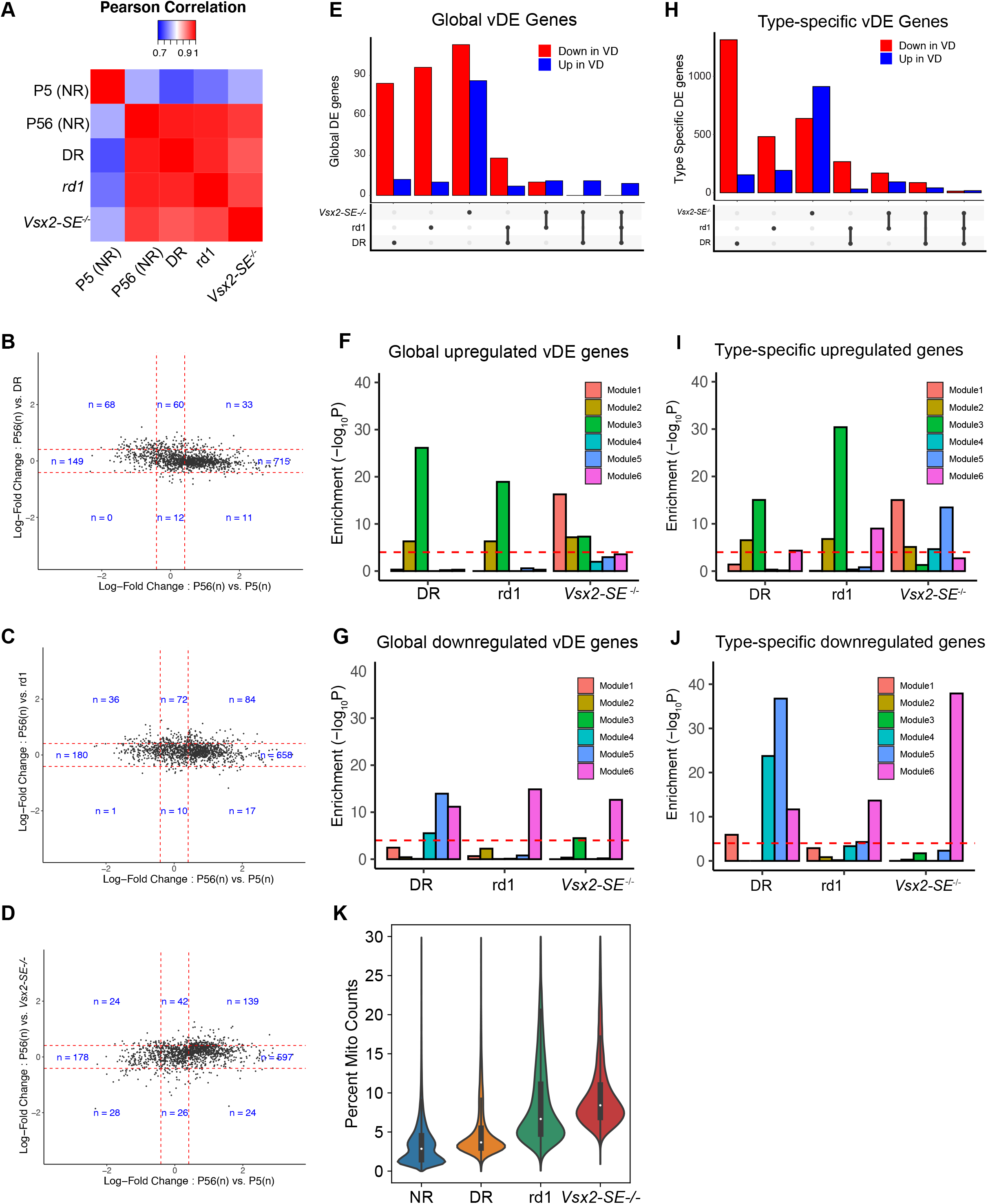
Global and type-specific gene expression changes in VD RGCs. A. Pairwise correlation heatmap showing that the average transcriptional profiles of VD RGCs are more similar to the NR RGCs at P56 than they are to NR RGCs at P5. Colors represent Pearson correlation coefficients. B. Scatter plot comparing average log2-fold changes in the expression of genes (points) between NR RGCs at P56 and NR RGCs at P5 (x-axis) versus between NR and DR RGCs at P56 (y-axis). Dashed red lines denote fold changes of 1.5 along each axis, and the number of genes in each region is indicated. The relative preponderance of dots with |log-fold| > 1.5 along the x-axis compared to the y-axis reflect the fact that maturation-related genes are not substantially impacted during VD. C. Same as B, for *rd1* RGCs. D. Same as B, for *Vsx2*-SE^-/-^ RGCs. E. Bar-plot showing the number of global vDE genes that are downregulated (red) or upregulated (blue) in a single VD condition compared to controls or shared among conditions. The combination of VD conditions corresponding to each pair of bars is indicated below. F. Statistical enrichment of maturation modules Mod1-6 (as in Figure 2A) among globally upregulated vDE genes in each VD condition. G. Same as F for globally downregulated vDE genes in each VD condition. H. Same as E but for type-specific vDE genes. I. Same as F but for type-specific upregulated vDE genes. J. Same as F but for type-specific downregulated vDE genes. K. Violin plot showing that the proportion of counts associated with mitochondrial transcripts is higher in the VD conditions compared to NR. Note that NR contains both control RGCs used in this study (Tran *et al.*, 2019) and RGCs from (Jacobi et al., 2022) (see Figure S5A).

In addition to global vDE genes, we also sought genes that were selectively upregulated or downregulated in each of the 45 types (type-specific vDE genes). We identified 3637 type-specific vDE genes that exhibited a >1.5-fold change (MAST DE test; adjusted p-value < 10^-4^), between at least one of the VD and the NR dataset in 5 or fewer types at a detection rate of ≥30% for either condition. We performed GO analysis on the combined set of global and type-specific vDE genes to assess pathways affected by VD (Figure S4A,B). We observed multiple instances of common GO terms enriched among vDE genes and those enriched in developmental modules. For example, most of the GO terms enriched in upregulated vDE genes in all three models were also enriched in developmental Mod2 and 3. However, it was challenging to interpret these similarities because of the redundancies among GO terms, a well-known problem (Jantzen et al., 2011). We therefore adopted the more direct approach of computing the statistical overlap between vDE genes and each of the six modules of temporally regulated genes identified in Figure 2. Upregulated genes in VD were significantly enriched for Mod 1-3, which are expressed in embryonic RGCs, while downregulated global vDE genes were enriched in Modules 4-6, which are postnatally active (p < 10^-4^, Hypergeometric test; Figures 7F,G). Similar trends were evident for type-specific vDE genes (Figures 7I,J). Together, these results suggest that visual deprivation directly impacts biological pathways involved in RGC development.

Remarkably, both global and type-specific vDE genes, were highly condition specific (Figures 7E,H). Some of these differences may result from the different ways in which the three models affected visual input, but it is also likely that some genes that are vDE in single models are false positives, resulting from inadequate sampling or technical variations among samples. Lacking additional replicates or an independent validation method, we therefore focused on groups of related genes. We highlight three interesting trends.

First, enrichment patterns of the modules were different among VD conditions: only in *Vsx2*-SE^-/-^ RGCs was Mod1 upregulated; only in DR mice were all three postnatal developmental modules downregulated; and only in *Vsx2*-SE^-/-^ were large numbers of genes upregulated (Figure 7F,G,I,J). Likewise, few GO enrichment terms were shared among conditions (Figure S4A,B).

Second, upregulated genes in *Vsx2*-SE^-/-^ RGCs included many implicated in the formation and function of excitatory postsynaptic specializations. GO terms included G-protein coupled receptor activity, regulation of postsynaptic density organization, postsynaptic density assembly, PDZ domain binding, and dendritic membrane (Figure S4). Upregulated genes included several encoding glutamate receptor subunits (*Gria1, Gria3, Grik3, Grik5*) as well as other components of excitatory postsynaptic densities (*Dlg1, Dlg4, Dlgap3, Lrrtm2, Lrrc4b, Ntrk3, Shank1*)(Holt et al., 2019). By preventing formation of bipolar cells, this mutant deprives RGCs of their main source of glutamatergic excitatory activation. The upregulation observed is reminiscent of “denervation supersensitivity” in which postsynaptic receptors and proteins associated with them are dramatically upregulated when skeletal muscle is denervated (Tintignac et al., 2015); similar phenomena have been observed in neurons (e.g.,(Kong et al., 2011; Kuffler et al., 1971)).

Third, mitochondrially-encoded genes were upregulated in all three VD models (Figure 7K and S5A). The upregulation was broadly shared among RGC types, being evident in all 45 types in rd1 and *Vsx2*-SE^-/-^ and in 36/45 types in DR (Figure S5B). Upregulated genes included *mt-Nd2*, *mt-Nd3*, *mt-Nd4*, *mt-Nd4l*, and *mt-Co3*, all of which have been found to bear missense (hypomorphic) mutations in Leber’s Hereditary Optic Neuropathy (LHON). Although these genes are ubiquitously expressed, the disease selectively affects RGCs (Yu-Wai-Man et al., 2011). Our results suggest the possibility that the decrease in mitochondrial gene expression caused by visual activity could further amplify respiratory chain dysfunction caused by the mutations, rendering RGCs particularly vulnerable to oxidative stress.

## DISCUSSION

The development of neurons, their differentiation into distinct types, and their integration into information processing circuits all result from hard-wired genetically encoded programs that are modified by neural activity. Both genetic and activity-dependent modes of development rely on molecular mediators, but our knowledge of their identities is incomplete for the former and rudimentary at best for the latter. Mouse RGCs are well suited for addressing these open questions for several reasons: (a) their structure, function and development have been studied in detail; (b) they comprise a neuronal class that has been divided into several subclasses and numerous (∼45) types, enabling analysis at multiple levels; and (c) methods are available for manipulating the sensory input they receive and thereby the patterns of activity they experience.

Our method was scRNA-seq, which enables comprehensive classification of neuronal cell types, and their mapping across developmental stages and experimental conditions. An additional advantage is that we were able to identify cells that were not RGCs and remove them from the dataset, ensuring that changes observed over time or after VD were attributable to RGCs and not to contaminating populations. By profiling RGCs at multiple developmental ages, we were able to map the changing transcriptional landscape of RGCs as they develop from embryonic stages to adulthood. By profiling adult RGCs that had been visually deprived in three different ways, we showed that vision is not required for full diversification of RGCs into subclasses and types but does affect patterns of gene expression in both global and type-specific ways. Our results can serve as a starting point for screening and assessing key molecular mediators of activity-independent and -dependent patterning of RGC development.

### RGC development

We first identified temporal gene expression changes broadly shared among developing RGC types. Both gene ontology and enrichment analysis of key gene classes (e.g., cell surface molecules, transcription factors, ion channels, neurotransmitter receptors) showed a systematic progression of expression patterns as RGCs differentiate, form synapses, and mature. DE genes expressed at E13 were enriched in TFs and regulators of neuronal differentiation and axon guidance. At later embryonic ages (E14 and E16), enriched genes included ones required for robust neuronal and axonal growth – for example, genes associated with ribosomal biogenesis and mitochondrial function. Perinatally (P0 and P5), genes required for synaptogenesis and synaptic choices – for example recognition molecules – are prominent, followed by genes encoding the machinery for axonal and synaptic signaling at P5 and P56 – for example, ion channels, neurotransmitter release components, and neurotransmitter receptors.

We next catalogued DE genes restricted to one or a few clusters at each age. TFs and CSMs were particularly prominent in this group. This is unsurprising in that genes involved in neuronal growth and function are shared among many neuronal types and classes. However, because the diversification into distinct RGC types occurs gradually, the relationship of embryonic clusters to adult types is not straightforward. We therefore used WOT, a statistical inference approach, to identify the likely precursors of each of the 45 types at each developmental stage. We used these assignments to trace the expression of two sets of TFs among these precursors: ones expressed in few precursor groups, which might be type-specific fate determinants, and ones previously suggested to be markers of RGC subclasses. Some potential type-specific TFs were expressed by characterized types for which reagents are available to test their roles – for example *Zic1* in W3B RGCs (C6), *Etv1* in alpha RGCs (C41, 42, 45; see also (Martersteck *et al.*, 2017)), and *Msc* in M2-ipRGCs (C31). Others are present in uncharacterized types and could be used to mark and manipulate them – for example *Eya1* in C10, *Runx1* in C13, and *Nfib* in C19 and 20. Of the TFs that defined subclasses, a few are expressed selectively at early times and might serve as fate determinants – for example, *Eomes/Tbr2*, *Mafb*, *Bnc2* and *Tfap2d*. Others are expressed selectively only peri- and postnatally – for example *Neurod2* and *Tbr1*. For those few that have been studied in retina, expression patters are consistent with mutant phenotypes: in *Eomes/Tbr2* mutants, most ipRGCs (a prominent set of *Eomes/Tbr2*-positive types) fail to develop, while in *Tbr1* mutants, T-RGCs develop but exhibit defects in dendritic morphogenesis (Kiyama *et al.*, 2019; Liu *et al.*, 2018; Sweeney et al., 2019).

Importantly, broader expression of other genes does not exclude the possibility that their functional roles may be type-specific. Specificity may arise due to combinatorial action of multiple genes, varying expression levels, or coupling with other molecular features. This may be particularly true for the broadly expressed TFs in Mod1 or the CSMs in Mod4 and Mod5 (Figure 2D; **Table S1**). Indeed, previous studies have found clear examples of redundancy among recognition molecules and TFs that regulate RGC development (e.g.(Duan *et al.*, 2018; Jiang *et al.*, 2013; Sajgo et al., 2017)).

### Visual deprivation

We used three methods to deprive mice of visual experience so we could ask how vision affects RGC diversification and maturation. Two methods relied on genetic manipulations – *rd1* mice in which photoreceptors degenerate beginning in the second postnatal week, and *Vsx2*-SE^-/-^ mice, in which no bipolar cells form, preventing communication from photoreceptors to RGCs. A third model, dark rearing from birth, prevents non-image forming vision noninvasively.

Importantly, although these perturbations prevent vision, they do not lead to complete inactivity of RGCs (Hooks and Chen, 2007); they therefore enabled us to assess roles of visually-evoked activity but not all electrical activity on RGC development. We observed that OFF RGCs in *Vsx2*-SE^-/-^ mice exhibited spontaneous bursty activity that was correlated between neighboring cells and was driven by inhibitory input likely arising from AII amacrine neurons. ON RGCs did not exhibit such spontaneous activity. This pattern resembles that previously described for RGCs in in *rd1* mice (Stasheff, 2008). Whether different patterns of spontaneous activity have a role in instructing type specific maturation is unclear. Unfortunately, although there are methods for disrupting the coherence of spontaneous activity among RGCs (Kirkby et al., 2013), it is currently not feasible to inhibit all action potentials in RGCs over a prolonged period. However, visual input has been shown to have clear effects on dendritic morphology of RGCs and refinement and maintenance of RGC axonal arbors in the superior colliculus (see **Introduction**), and transcriptomic analyses have documented significant alterations in gene expression in the lateral geniculate nucleus and visual cortex following visual deprivation (Cheadle et al., 2018; Cheng et al., 2022; Hrvatin et al., 2018; Tropea et al., 2006).

The observation of fewer RGC clusters in VD retina than in NR retina initially suggested that late steps in diversification of types might require visual input. However, when assessed at a cell-by-cell level, over 98% of RGCs mapped with high confidence to a single adult type, even in clusters that contained RGCs of two or three types. Further, in terms of transcriptomic similarity, VD RGCs unequivocally resembled aged matched normally reared counterparts, rather than immature RGCs at P5. Previously, we have shown that RGC diversification is incomplete at P5 (Shekhar *et al.*, 2022). The present results suggest that non-image forming visual-activity during the early postnatal period is not required for the establishment of RGC diversity, and image-forming light-driven activity following eye opening is not required for its maintenance. However, we cannot rule out the possibility that light-independent activity, which begins by E16, influences RGC diversification.

VD had clear effects on gene expression in RGC generally as well as in specific RGC types. In aggregate, they suggested maturational defects. First, VD RGC types exhibited fewer DE genes than their NR counterparts. Second, the genes that were altered were enriched for those that are temporally regulated during normal development. Genes expressed at embryonic stages of RGC development and subsequently downregulated (Mods1-3) were expressed at higher levels in VD RGCs than normal RGCs, while genes upregulated in postnatal RGCs (Mods4-6) were expressed at lower levels in VD RGCs. We speculate that these subtle but systematic changes may be associated with previous studies showing that dark rearing perturbs both dendritic and axonal refinement (see above), without leading to a complete dedifferentiation of RGCs.

In all three VD models, we profiled adult RGCs. A remaining question is whether vision affects maturation per se, maintenance of mature characteristics, or both. Physiological studies have provided evidence for both possibilities (e.g., (Carrasco et al., 2005; Feldheim and O’Leary, 2010; Hooks and Chen, 2006; Hooks and Chen, 2007). Profiling of RGCs at earlier times could settle this issue.

## DATA AVAILABILITY STATEMENT

The gene expression matrices and analysis scripts are available on our lab’s Github repository at https://github.com/shekharlab/RGC_VD. Requests may be directed to Karthik Shekhar at.

## AUTHOR CONTRIBUTIONS

I.E.W, J.R.S. and K.S. conceived the project and designed the study; I.E.W. performed the single-cell RNA-seq and histology experiments; F.R. performed the physiology experiments; S.B. and designed the computational methods and performed the data analysis; M.D. provided the *Vsx2*-SE^-/-^ line prior to publication; S.B., J.R.S and K.S. wrote the paper with input from all authors.

---

**Figure S1.**
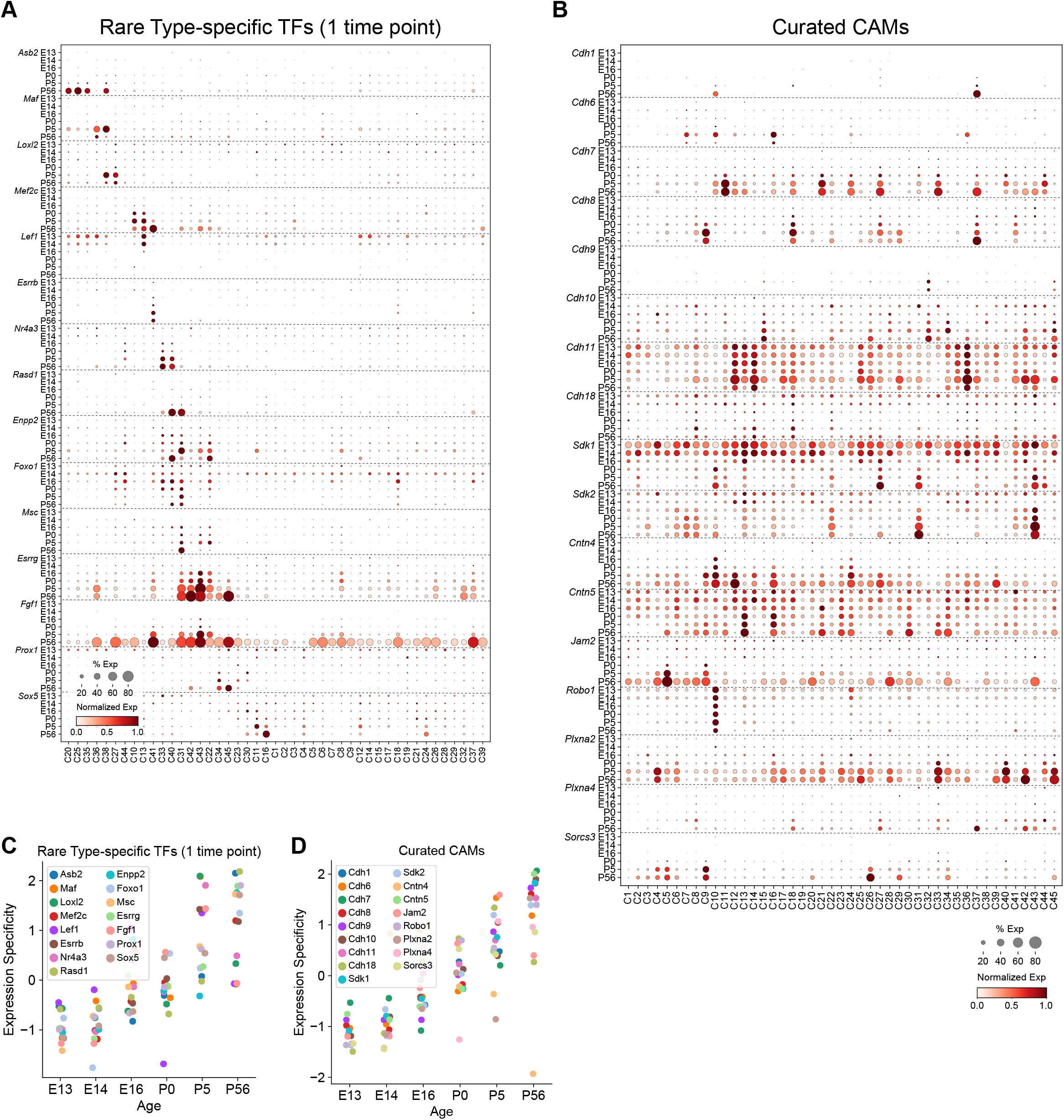
Rare type-specific TFs and assessment of CSM expression. A. Dot plot showing expression patterns of type-specific TFs (rows) with age in putative type-specific precursors (columns). Representation as in Figure 4D. This panel only displays TFs that were identified as type-specific in at least one of the six ages. B. Same as panel A, displaying expression patterns of a subset of CSMs known to play roles in synaptic choices of RGCs (see **Results**). C. Expression specificity type-specific TFs shown in panel A with age. Unlike TFs in Figure 4F, here certain TFs become less specific with age. D. Expression specificity with age for CSMs in panel B.

**Figure S2.**
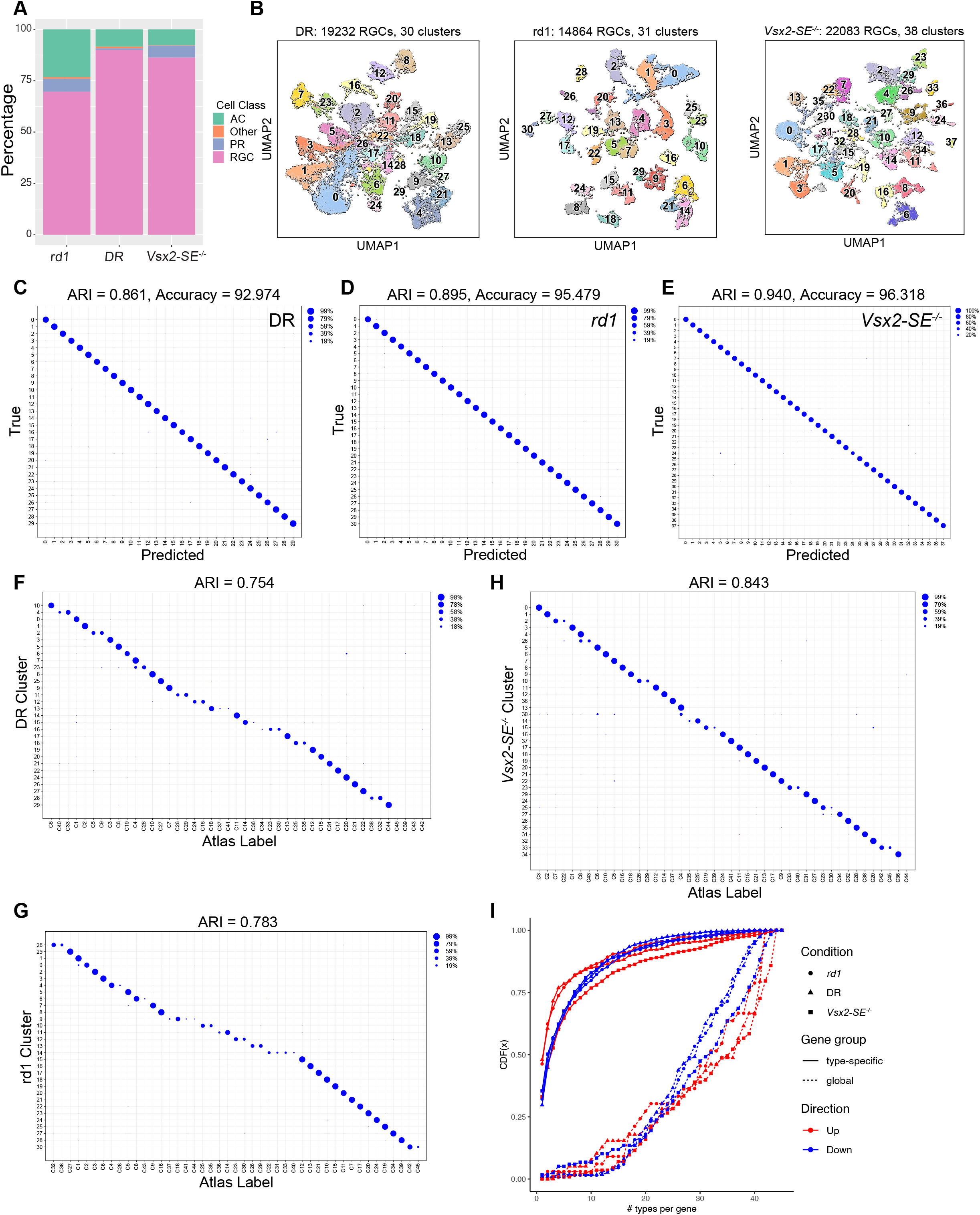
Analysis of scRNA-seq data from three visual deprivation conditions. A. Composition (y-axis) of major cell classes in the scRNA-seq datasets collected from three visually deprived experiments (x-axis). RGCs comprised a majority (>70%) in all collections, while the main non-RGC classes included ACs and PRs. The minority category labeled as “other” predominantly comprised immune and endothelial cells. B. Left to right: Panels showing UMAP visualization of DR RGCs, *rd1* RGCs and *Vsx2*-SE^-/-^ RGCs colored by cluster identity. The coordinates are identical to Figures 6B-D. C. Confusion matrix showing that all transcriptomically defined clusters in DR RGCs as in panel B are learnable by a classifier with >90% accuracy. ARI: adjusted Rand index, a measure (−1 to 1) of mapping specificity. D. Same as C for *rd1* RGCs. E. Same as C for *Vsx2*-SE^-/-^ RGCs. F. Confusion matrix showing mapping between DR clusters and NR types at P56 as in Tran et al., 2019. G. Same as F, for *rd1* RGCs H. Same as F, for *Vsx2*-SE^-/-^ RGCs. I. Post-hoc consistency check for global and type-specific vDE genes in each condition (shape). Each of the 12 curves corresponds to a set of vDE genes identified in a condition (*rd1*, DR, or Vsx2-SE^-/-^), DE category (global or type-specific) and DE direction (up or down with respect to NR RGCs). Each vDE curve plots the cumulative distribution function or CDF (y-axis) corresponding to the number of types that show a significant change. CDF(x) denotes the fraction of genes in the vDE set that are DE in < x types. Thus, the “convex up” shape of the global vDE CDF suggests that these gene alterations are broadly shared. In contrast, the “concave up” shape of the type-specific CDFs suggests that these alterations are present only in a subset of types.

**Figure S3.**
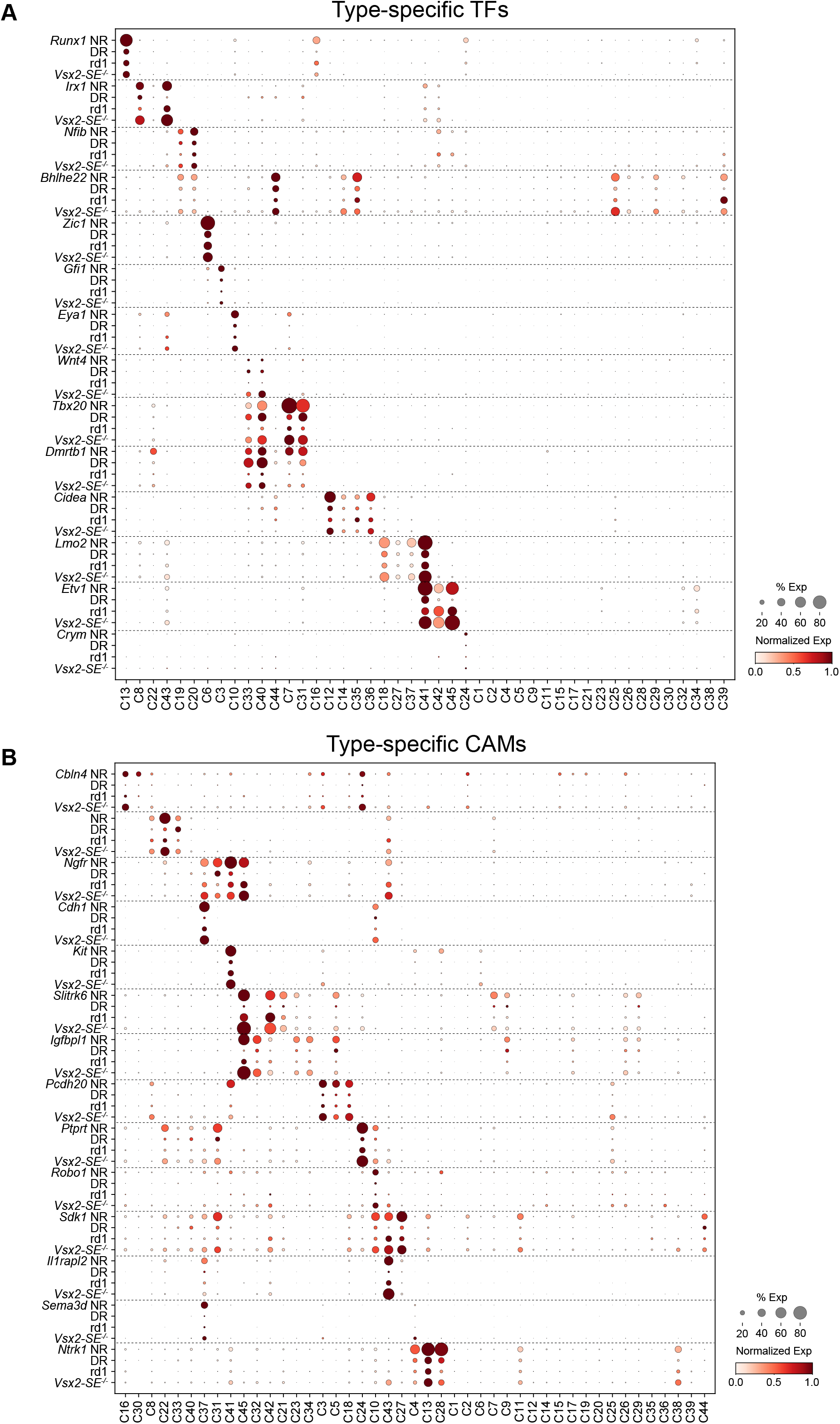
Type-specific gene expression of TFs and CSMs is maintained in VD. A. Dot plot showing expression patterns of type-specific TFs in VD retina. Developmental profiles are shown in Figure 4E. The size of each dot corresponds to the fraction of cells with non-zero transcripts, and color indicates normalized expression levels. Row blocks corresponding to different TF are demarcated by dotted horizontal lines. B. Same as A, but for a subset of the CSMs, including some shown in Figure S1B.

**Figure S4.**
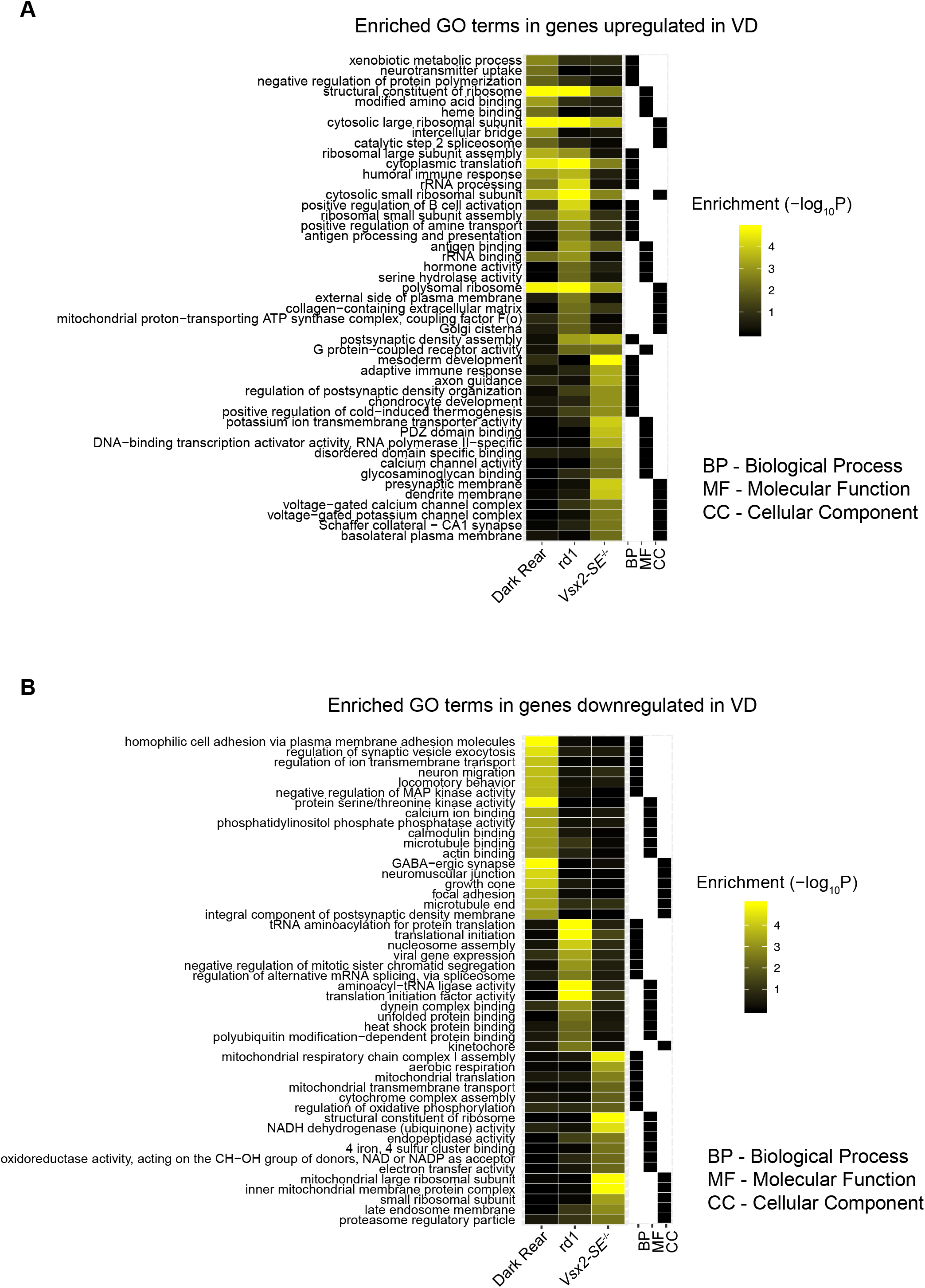
Analysis of vDE genes. A. Heatmap displaying significant GO terms (rows) associated with genes (global and type-specific combined) that are upregulated in VD (columns) compared to NR RGCs. Values correspond to −log10(adjusted p-value). Black and white annotation bar on the right of the heatmap identifies the GO category corresponding to each row (BP, MF or CC). B. Same as A, for downregulated genes in VD samples compared to control.

**Figure S5.**
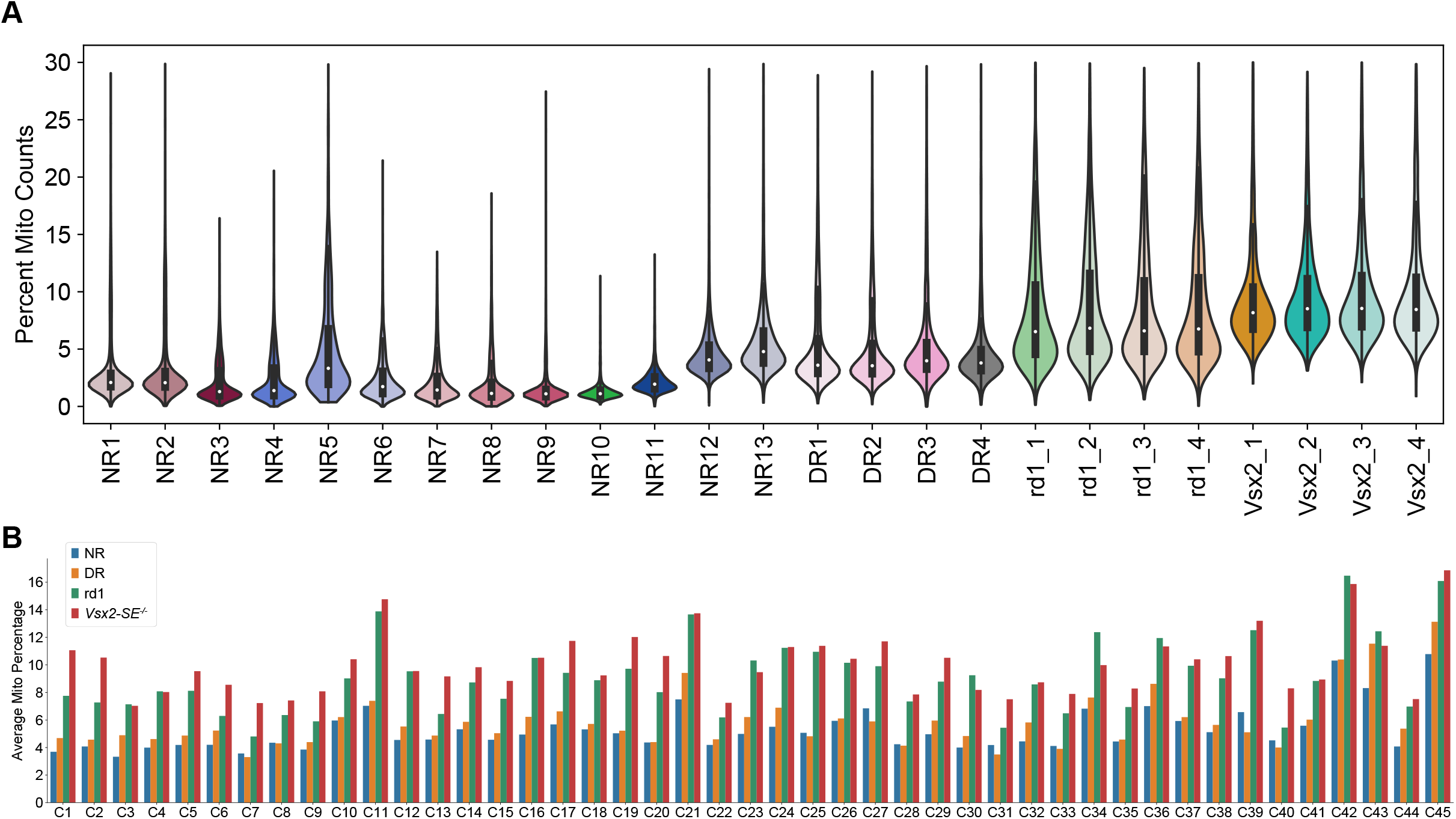
Expression of mitochondrially encoded genes in RGC types and conditions. Violin plot showing that the percentage of counts associated with mitochondrial transcripts is higher in the VD biological replicates compared to NR. NR1-10 are from (Jacobi *et al.*, 2022) and constitute 36,620 RGCs not used in this study except in **Figure 7K**. NR11-13 are from (Tran *et al.*, 2019) and constitute the 35,699 NR RGCs used throughout this study in comparative analyses to VD RGCs. Batches were analyzed separately and the additional control data were included because apparent increases in mitochondrial gene expression are sometimes observed when cells are damaged during processing (Stegle et al., 2015). The consistent differences between NR and VD samples provides evidence that batch-to-batch variation does not account for the upregulation of mitochondrial gene expression we observed. B. Bar plot showing the mean proportion of mitochondrial transcripts in each RGC type across all four conditions. The upregulation of mitochondrial genes in VD shows no apparent type specificity.

## SUPPLEMENTARY TABLE LEGENDS

**Table S1. Global modules of temporally regulated genes during RGC development.**

**Table S2. Cluster-specific genes at each age during RGC development.**

**Table S3. Global vDE genes upregulated in two out of three VD conditions.**

## Acknowledgments

Funding for this study was provided by National Institute of Health grants R37NS029169, R01EY022073 (JRS), and R00EY028625 (KS), NSF GRFP DGE1752814 (SB), the Chan-Zuckerberg Initiative (JRS, KS) and startup funding from UC Berkeley (KS, SB). The authors would like to thank Drs. Chinfei Chen and Marla Feller for helpful discussions.

